# Personalized Neoantigen Vaccines Synergize with Immune Checkpoint Therapy and CD8-Targeted Cytokines to Control B-Cell Lymphoma

**DOI:** 10.64898/2026.08.02.742304

**Authors:** Yuang Song, Ekaterina Aladyeva, Ruan F. Vieira Medrano, Derek J Theisen, Cora D. Arthur, J Michael White, Heather Brink Kohlmiller, Anthony Vomund, Kartik Singhal, My Hoang, Samuel Ameh, Kathleen C.F. Sheehan, Ronald Levy, Todd A. Fehniger, Maxim N. Artyomov, Malachi Griffith, Obi L. Griffith, Yik Andy Yeung, Ivana Djuretic, Hussein Sultan, Robert D. Schreiber

**Author notes:** **Co-Corresponding Authors Information:** Dr. Hussein Sultan, Dr. Robert D. Schreiber.

## Abstract

Personalized neoantigen (neoAg) vaccines have shown clinical promise in solid tumors^1–8^, yet their efficacy and mechanism of action in hematopoietic malignancies remain poorly defined^9–11^. Herein, we establish an immunocompetent syngeneic A20 B-cell lymphoma platform to test the efficacy of neoAg vaccines used either as mono- or combinatorial therapies with other immunotherapies^12–17^. Whereas subcutaneous A20 tumors were refractory to single-agent αPD-1 or αCTLA4 therapy, they were eradicated in a T cell-dependent manner in 90% of syngeneic hosts treated with dual immune checkpoint therapy (dual ICT, i.e., αPD-1 + αCTLA4). By mapping antigen specificity of dual-ICT-elicited T cells, we identified and validated dominant endogenous A20 MHC-I and MHC-II neoantigens and designed therapeutic synthetic long peptide (SLP) vaccines containing these neoepitopes. This vaccine (A20 neoVAX) promoted robust neoAg-specific CD4□ and CD8□ T cell responses in naïve syngeneic BALB/c mice and induced tumor rejection in ∼70% of subcutaneous tumor-bearing mice. In addition, nearly all mice rejected their subcutaneous A20 tumors when A20 neoVAX was combined with αPD-1. To render the results of this study more physiologic, we developed a systemic A20 lymphoma model and found that dual ICT failed to control tumor progression and A20 neoVAX delayed tumor progression and prolonged animal survival but did not induce tumor rejection. In contrast, A20 neoVAX plus dual ICT achieved durable systemic tumor elimination. Mechanistically, the combination of A20 neoVAX plus dual ICT amplified priming of A20 neoAg-specific T cells, prevented T cell dysfunction, sustained the cytotoxic capacity of tumor-specific CD8^+^ T cells, and induced Th1-skewing of CD4^+^ T cells in tumor and peripheral compartments. To increase the clinical relevance of these findings and to minimize potential adverse events in tumor-bearing, therapeutically treated individuals, we substituted CD8-targeted cytokine muteins (CD8-IL2 or CD8-IL21) for αCTLA4. These agents represent genetically modified forms of IL-2 or IL-21 that selectively stimulate CD8^+^ T cells but have significantly reduced capacity to activate chronic inflammation and immunosuppressive functions of other immune cells. Whereas mice bearing systemic A20 lymphoma treated with either nothing, A20 neoVAX, or A20 neoVAX + CD8-IL2 failed to control tumor outgrowth, 66.7% of tumor-bearing mice treated with A20 neoVAX + CD8-IL2 + αPD-1 rejected their tumors. In similar experiments in which CD8-IL21 was substituted for CD8-IL2, tumor clearance was also observed in two-thirds of A20-bearing mice but now rejection occurred in the absence of αPD1. Together, these data define a framework for optimal personalized neoAg vaccination in B-lymphoma and demonstrate that neoAg vaccines can safely synergize with CD8^+^ T cell-selective immunotherapies to prevent T-cell dysfunction and generate durable systemic anti-tumor immunity.

## INTRODUCTION

Immune checkpoint therapy (ICT) has transformed cancer treatment by reversing T-cell dysfunction and enhancing anti-tumor immunity^18,19^. Recent clinical and mechanistic studies indicate that responses to checkpoint inhibitors strongly depend on endogenous immunity against neoantigens (neoAg), suggesting that augmenting such responses may improve clinical outcomes^20–23^. Therapeutic cancer vaccines, capable of priming and expanding neoAg-specific CD8□ and CD4□ T cells, provide therapeutic potential when used as monotherapy and offer a rational complementary strategy to enhance ICT efficacy^5,24–26^.

Personalized neoAg cancer vaccines have shown encouraging immunologic and clinical activity in patients with solid tumors^1–8^. Platforms ranging from synthetic long peptides to mRNA, DNA, and dendritic cell vaccines have induced detectable circulating and intratumoral neoAg-specific T cells, accompanied by delayed or absent tumor relapse and evidence of immunologic response^1–8^. Furthermore, the combination of neoAg cancer vaccines plus ICT has proven feasible and well-tolerated in early-phase clinical trials^3,4^. Multiple studies have demonstrated functional synergy between the two modalities, characterized by enhanced T cell priming, mitigation of exhaustion phenotypes, and improved tumor control.

Despite rapid expansion of evidence in solid cancers, the therapeutic potential of neoAg vaccines for hematologic malignancies such as B-cell lymphomas remains insufficiently studied^9–11^. B-cell lymphomas present a distinct immunologic landscape, including altered antigen presentation, altered T-cell infiltrates, and immune evasion within lymphoid tissues^27–31^. While mutation burdens may be lower relative to certain solid tumors^32^, lymphoma genomes frequently harbor mutation-derived epitopes arising from somatic hypermutation, class switch recombination, and other oncogenic processes, suggesting that neoAg targeting may be feasible^9,10^. Indeed, in a small pilot clinical trial of personalized neoAg vaccines plus anti-PD1, clinical responses and neoAg-specific T cells were observed^10^. However, extensive systematic evaluation of neoAg vaccine efficacy, T cell responses, and interactions with the lymphoma microenvironment in immunocompetent models remains lacking.

In this study, we evaluated the therapeutic efficacy and immunologic mechanisms of action of neoAg vaccines in syngeneic mouse models of B-cell lymphoma. Employing the widely used A20 lymphoma model in BALB/c mice^12–17^, we characterized neoAgs arising from A20-expressed mutations and designed A20-specific neoantigen therapeutic synthetic long peptide (SLP) vaccines (A20 neoVAX). We assessed A20 neoVAX-induced tumor control, immune microenvironmental remodeling, and functional interactions with checkpoint pathways and cytokine signaling in both subcutaneous and systemic A20 lymphoma models. We demonstrate a difference in response to therapy between the subcutaneous and systemic models and document that combinatorial vaccine-based immunotherapies are required for durable tumor control in the more physiologic systemic lymphoma model. We also show that effective vaccine-based therapies promote tumor rejection by circumventing T-cell dysfunction. These findings thus demonstrate the therapeutic potential for neoantigen vaccines in hematologic malignancies, provide translational strategies for rational combination immunotherapies, and suggest how personalized cancer vaccines may be effectively used for the treatment of B lymphoma.

## RESULTS

### Dual ICT induces A20 neoAg-specific T cells and leads to rejection of subcutaneous A20 lymphoma

The subcutaneous A20 lymphoma model has been widely used to study B-cell lymphoma responses to chemotherapies and immunotherapies^13–17^. Previous reports indicate that subcutaneous A20 is resistant to single-agent ICT such as αCTLA4^13–15,17^, reflecting the limited clinical responsiveness of many non-Hodgkin lymphomas (NHL)^33–35^. To assess the immunogenicity of A20 lymphoma in vivo and characterize endogenous immune responses elicited by ICT, we first evaluated tumor control following single or dual ICT in immunocompetent BALB/c mice bearing subcutaneous A20 tumors **(Fig. 1a)**. While treatment with either αPD-1 or αCTLA4 induced limited tumor rejection in 20-26% of mice bearing subcutaneous A20 tumors, dual ICT with αPD-1 plus αCTLA4 induced tumor rejection in 90% of treated mice **(Fig. 1b, c)**.

**Figure 1:**
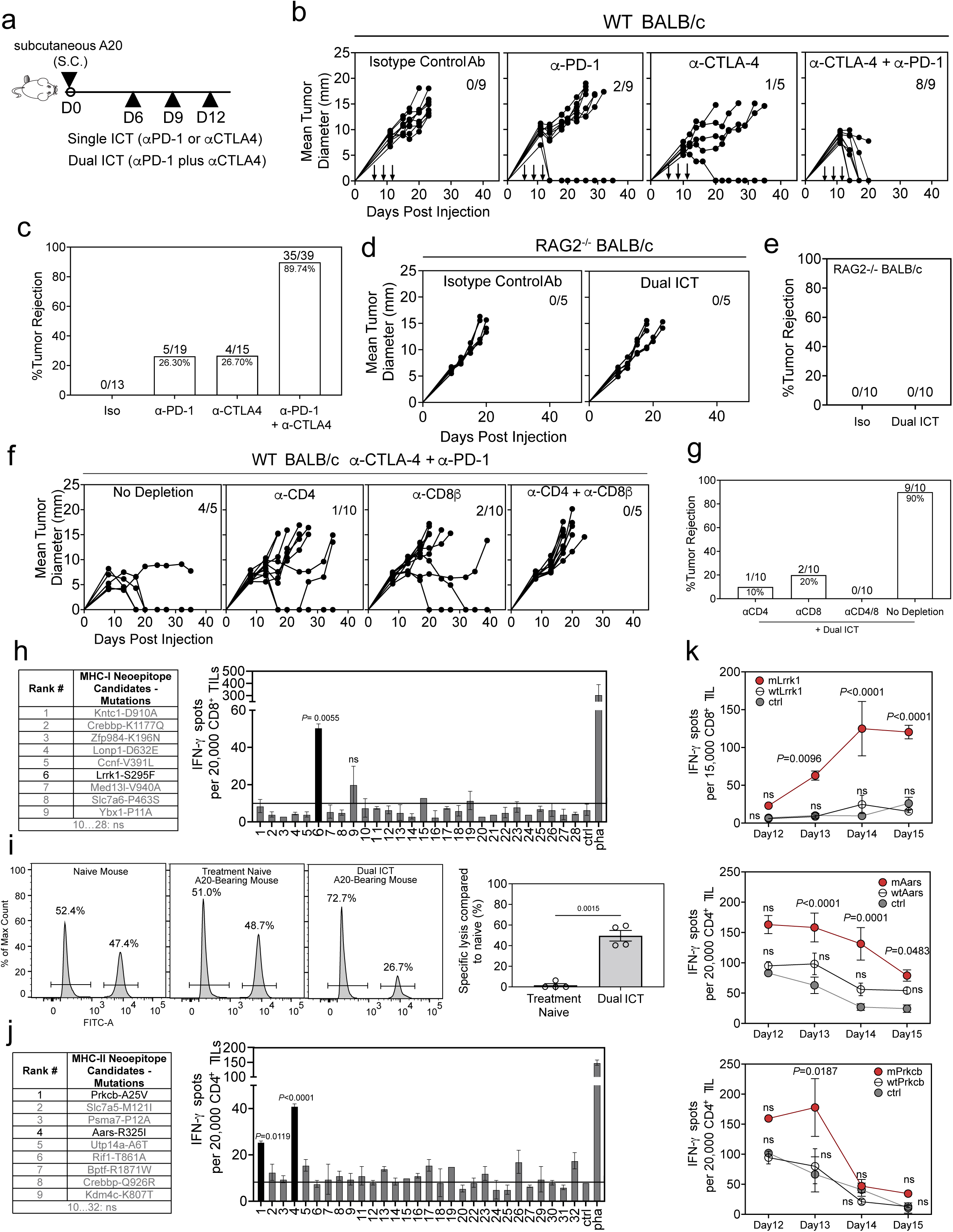
Immune checkpoint blockade mediates rejection of subcutaneous A20 lymphoma and enables identification of immunogenic neoantigens. **a,** Schematic of the ICT treatment schedule. **b,** Representative A20 tumor outgrowth over time (numbers in the panels indicate the number of mice that rejected their tumors versus the total number of mice included in this experiment). **c,** Percent survival of mice receiving the indicated treatments across multiple replicate experiments. Combined results from 3 studies conducted as in panel b are reported as the number of mice that rejected their tumors divided by the total number of mice used in replicate experiments. **d,e** Representative A20 tumor outgrowth in RAG2□/□ BALB/c mice following control mAb or dual ICT treatment (d), and overall percent survival across several experimental replicates, **(e). f,** Representative A20 tumor outgrowth and **g,** percent survival following immune checkpoint blockade with or without depletion of CD4□ T cells, CD8□ T cells, or both. Exact sample sizes are reported as the number of mice that rejected tumors divided by the total number of mice used in the experiment. **h, (left)** predicted MHC-I-restricted neoantigen peptides used to stimulate tumor-infiltrating sorted CD8□ T cells from dual ICT-treated tumor-bearing mice, **(right)** results of IFNγ ELISPOT analyses. Data are shown as mean □±□ s.e.m. from n = 3 biological replicates (indicated groups compared with control group, one-way ANOVA with Dunnett multiple comparisons test). **i,** Flow cytometry-based in vivo cytotoxicity assay. **Left,** representative histogram. **Right,** quantification of target cell killing by effector cells isolated from untreated or ICT-treated tumor-bearing mice compared to naïve mice. Quantification shows mean □±□ s.e.m. from n = 4 biological replicates, repeated two times (Welch’s t-test). **j, (left)** predicted MHC-II-restricted neoantigen peptides used to stimulate tumor-infiltrating CD4□ T cells sorted from dual ICT-treated tumor-bearing mice, **(right)** results of IFNγ ELISPOT analyses. Data are shown as mean □±□ s.e.m. from n = 3 biological replicates (indicated groups compared with control group, one-way ANOVA with Dunnett multiple comparisons test). k, IFNγ ELISPOT analysis of CD8□ TILs **(top)** following stimulation with mutant or wild-type Lrrk1 peptides, and CD4□ TILs following stimulation with mutant or wild-type Aars **(middle)** or Prkcb **(bottom)** peptides. Data are shown as mean □±□ s.e.m. from n = 4 biological replicates, repeated two times (P values are noted in figures representing indicated groups compared with the control group, two-way ANOVA with Tukey multiple comparisons test).

To determine whether adaptive immunity was required for dual ICT-mediated tumor rejection, we tested the therapeutic antitumor efficacy of dual ICT in BALB/c RAG2 □^/^□ mice lacking T cells, B cells, and NKT cells or in WT mice after depleting CD4^+^ or CD8^+^ T cells either alone or together. Dual ICT failed to control subcutaneous A20 tumor growth in BALB/c RAG2 □^/^□ mice, demonstrating a requirement for adaptive immune responses **(Fig. 1d, e)**. Monoclonal antibody depletion of either CD4□ or CD8□ T cells alone or together in WT mice also abrogated subcutaneous A20 rejection following dual ICT **(Fig. 1f, g)**. Consistent with these findings, treatment with dual ICT was associated with increased spatial proximity between CD4□ and CD8□ tumor-infiltrating lymphocytes (TILs) and enhanced interactions between lymphocytes and dendritic cells within the tumor microenvironment (TME) similar to our previous findings using the T3 methylcholanthrene-induced sarcoma model^8,21,22,36,37^ **(Supplementary Fig. 1a)**.

Given the requirement for both CD4□ and CD8□ T cells in dual ICT-induced tumor rejection, we next investigated whether ICT-promoted immunity to tumor-specific neoAgs. Whole-exome and RNA sequencing of A20 cells versus normal cells identified 233 expressed nonsynonymous somatic mutations, a number that is comparable to the A20 mutational load reported by others^38^. Using the hmMHC and pVACtools algorithms reported previously^22,39–42^, we predicted candidate neoAg peptides derived from mutant A20 proteins with binding potential to MHC-I (H2-K^d^, H2-D^d^, H2-L^d^) or MHC-II (I-A^d^, I-E^d^) alleles.

To functionally assess antigen recognition, we performed IFNγ ELISPOT assays using tumor-infiltrating CD4^+^ or CD8□ T cells isolated from dual ICT-treated subcutaneous tumor-bearing mice. This screening identified a limited number of immunogenic mutations recognized by CD4□ and CD8□ TILs. Among MHC-I-restricted candidates, CD8□ TIL robustly responded to a major mutant H2-L^d^-restricted peptide derived from leucine-rich repeat kinase 1 (Lrrk1: ENSMUST00000015277) containing an S295F substitution (mLrrk1), but not the corresponding wild-type peptide **(Fig. 1h, Supplementary Fig. 1b)**. By screening truncated mLrrk1 peptides (SLP, 25 amino acids), we identified the minimum 9 amino acid MHC-I binding core (LPFIIPWGL) from mLrrk1 **(Supplementary Fig. 1c)**. We then employed this minimal sequence to generate a mLrrk1-H2-L^d^ tetramer and showed that it stained subsets of CD8^+^ TILs isolated from subcutaneous A20 tumors **(Supplementary Fig. 1d)**. In vivo cytotoxicity assays further demonstrated that CD8□ T cells induced by dual ICT specifically lysed target cells presenting mLrrk1 peptide, confirming functional neoantigen-specific cytotoxicity **(Fig. 1i)**. Thus, our independent data confirm the identification of a major A20 MHC-I-restricted antigen reported by the J. Brody lab^43^.

Similarly, ELISPOT screening of MHC-II-restricted candidates identified CD4□ T cell responses specific to an R325I mutation in alanyl-tRNA synthetase 1 (mAars, Aars: ENSMUST00000034441) and an A25V mutation in protein kinase C beta (mPrkcb, Prkcb: ENSMUST00000064989) (Fig. 1j). Both antigens were H-2 I-E^d^ restricted. Reactivity was preferentially directed against mutant peptides and was minimal or absent against the corresponding wild-type sequences **(Supplementary Fig. 1b)**. Using approaches similar to those employed for MHC-I neoAgs, we generated mPrkcb-I-E^d^ tetramers that bound to mPrkcb-specific CD4^+^ T cells **(Supplementary Fig. 1e)** and showed that they only stained CD4^+^ T cells from mice bearing subcutaneous A20 tumors but not CD4^+^ T cells from normal control mice.

We next examined the temporal dynamics of neoantigen-specific T cell responses following dual ICT. IFNγ ELISPOT analysis revealed that neoantigen-specific CD4□ T cells were maximally detected in the tumor between days 12 and 13 and preceded the maximal appearance of intratumoral neoAg-specific CD8^+^ T cells by one day i.e., day 14 **(Fig. 1k)**. These data indicate distinct kinetics of CD8□ and CD4□ T cell responses during ICT-mediated rejection of subcutaneous A20 tumors.

Together, these results demonstrate that dual αPD-1 and αCTLA4 treatment induces coordinated A20-specific CD4□ and CD8□ T cell responses capable of driving efficient rejection of subcutaneous A20 lymphoma.

### An A20-specific NeoAg SLP vaccine (A20 neoVAX) can eliminate established subcutaneous A20 lymphomas in vivo

Previous studies have shown that SLPs encoding neoAg epitopes can be used to design therapeutic vaccines that elicit T cell-mediated tumor rejection in solid tumor models^1,22,24,25^. To assess the immunogenicity of A20-derived MHC-I and MHC-II neoAgs delivered in SLP vaccines, naïve BALB/c mice were vaccinated with 25-mer SLPs encoding the MHC-I-restricted neoAg mLrrk1 and the MHC-II-restricted neoAgs mAars and mPrkcb, formulated with poly-ICLC adjuvant. IFNγ ELISPOT analysis of splenocytes from vaccinated mice demonstrated preferential recognition of mutant peptides, with minimal reactivity to corresponding wild-type or control peptides, confirming that SLP vaccination induced neoAg-specific T cell responses **(Supplementary Fig. 2a-d)**. These data confirm that both MHC-I- and MHC-II-restricted A20 mutations are immunogenic when delivered as SLP vaccines.

We next studied the activity of A20 MHC-I and MHC-II neoAg responses in promoting antitumor efficacy **(Fig. 2a)**. Vaccination with an irrelevant control SLP failed to affect tumor growth, and vaccines containing either the MHC-I (mLrrk1) or MHC-II (mPrkcb) A20 neoAg alone conferred only minimal therapeutic benefit **(Fig. 2b, c)**. In contrast, therapeutic vaccination of mice bearing subcutaneous A20 tumors with a combination of MHC-I and MHC-II A20 neoAgs (A20 neoVAX) resulted in efficient rejection of established tumors in 23/33 mice **(69.7%, Fig. 2b, c)**. These findings point to a need for both CD4^+^ and CD8^+^ T cells to achieve tumor elimination. Switching the MHC-II neoAg mPrkcb to the other A20 MHC-II neoAg mAars also produced a similar therapeutic efficacy of 19/28 (67.9%) responding mice, **Supplementary Fig. 2e**), indicating a reproducible therapeutic benefit of dual-compartment neoAg targeting. Additionally, vaccines containing the MHC-I-restricted neoAg together with either only one or both MHC-II-restricted A20 neoAgs exhibited equivalent antitumor efficacy, resulting in comparable tumor rejection **(Supplementary Fig. 2f)**. These findings document that MHC-I neoAg responses alone are insufficient for tumor rejection and highlight the requirement for MHC-II neoAg responses to achieve effective tumor control in the A20 B-cell lymphoma model, consistent with previous studies from our lab and others^22,24,44,45^.

**Figure 2:**
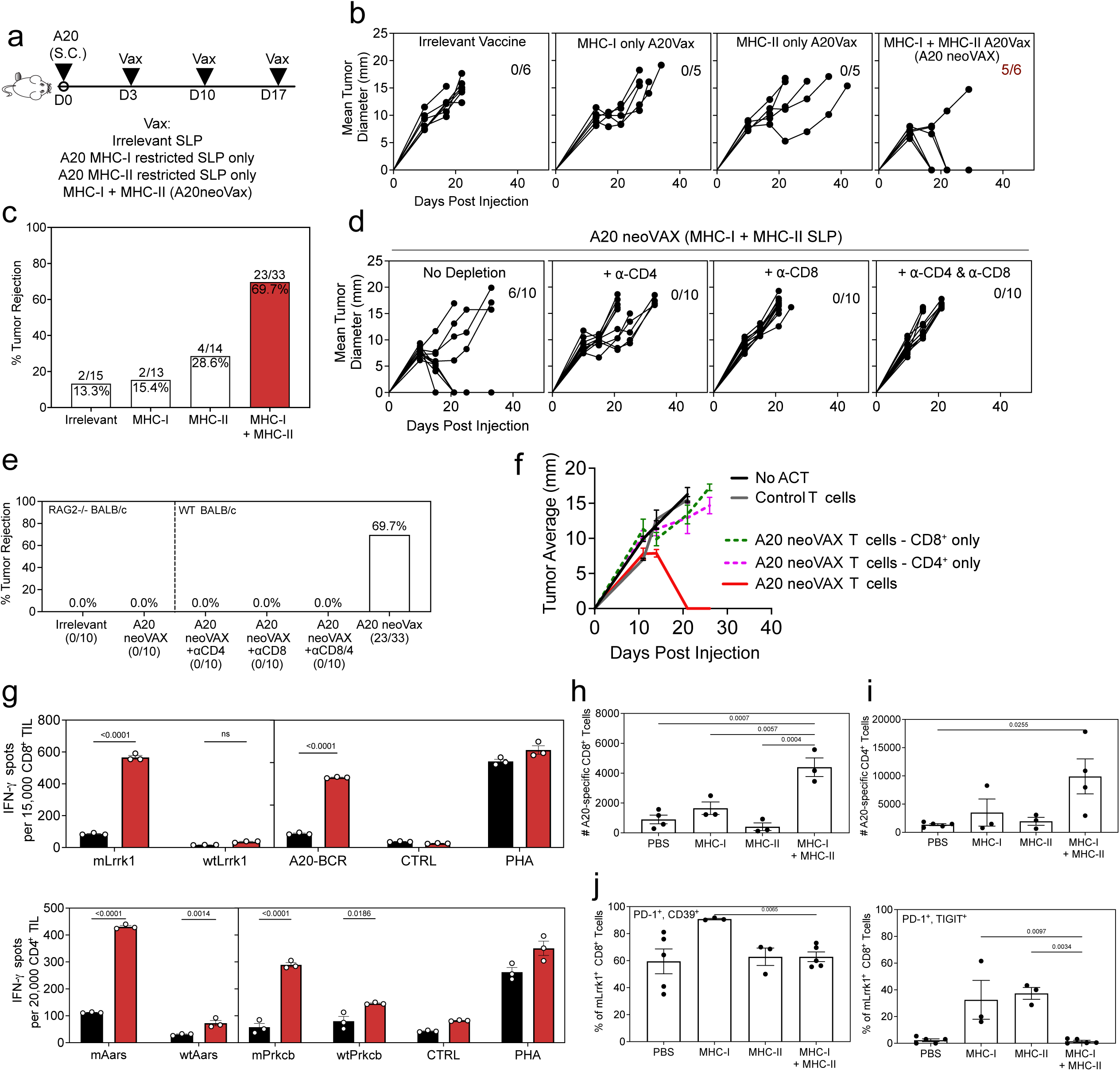
Neoantigen peptide vaccination induces coordinated CD8□ and CD4□ T cell-dependent control of subcutaneous A20 lymphoma. **a,** Schematic of the vaccination schedule and treatment groups. **b,** A20 tumor outgrowth curves following vaccination with different derivatives of A20 vaccines. Numbers indicate the number of mice in each group that rejected their tumors (complete responders) versus the total number of mice in the group. **c,** Percent survival of mice receiving the indicated vaccines, shown as the number of mice that rejected their tumors over total mice per group (combined from 5 independent experiments for the MHC-I+MHC-II group; or 3 experiments for other groups). **d,** Tumor growth curves following therapeutic vaccination with A20 neoVAX (combined MHC-I + MHC-II neoantigen SLP) in WT mice or following depletion of CD8□ T cells, CD4□ T cells, or both. **e,** Tumor rejection frequencies in WT or RAG2□/□ BALB/c mice, following neoantigen vaccination with or without T cell depletion. **f,** Tumor outgrowth in RAG2□/□ BALB/c mice following adoptive transfer of vaccine-induced CD8 T cells, CD4 T cells, or the combination of both. A20 neoVAX was used to generate donor T cells. Individual tumor growth curves were compared with tumor growth in recipients receiving total vaccine-induced T cells using two-way ANOVA (mixed-effects model, Tukey multiple-comparison test). **g,** IFNγ ELISPOT analysis of neoantigen-specific CD8□ TILs **(top)** and CD4□ TILs **(bottom)** from control vaccine or A20 neoVAX-treated mice (n=3, two-way ANOVA Šídák’s multiple comparisons test). **h,** Quantification of numbers of neoantigen-specific CD8^+^ TILs (n=4 for PBS-treated group, n=3 for other groups) and **i,** CD4^+^ TILs, (n=5 for PBS-treated group, n=4 for MHC-I+MHC-II-treated group, n=3 for other groups). **j,** Expression of inhibitory receptors PD-1 and CD39 **(left)** or PD-1 and TIGIT **(right)** on mLrrk1-specific CD8□ T cells following vaccination with PBS, MHC-I, MHC-II, or combined MHC-I + MHC-II neoantigen vaccines (n=5 for PBS/MHC-I+II-treated groups, n=3 for other groups). Statistical analysis for h-j was performed using one-way ANOVA with Tukey multiple comparison test. Data are shown as mean □±□ s.e.m.

To more precisely define the immune cell populations required for vaccine-mediated tumor rejection, CD4□ and/or CD8□ T cells were eliminated using depleting subset-specific mAbs during A20 neoAg vaccination. Depletion of either subset or combined depletion eliminated the vaccine’s therapeutic efficacy **(Fig. 2d, e)**. This result demonstrates that both CD4□ and CD8□ T cells are required for vaccine-induced tumor rejection. In a complementary approach, adoptive cell transfer experiments demonstrated that transfer of vaccine-induced T cells into immunodeficient, tumor-bearing recipients conferred tumor control, whereas transfer of control T cells did not **(Fig. 2f)**. Transfer of purified CD4□ or CD8□ T cells alone resulted in reduced efficacy compared with transfer of both subsets together, supporting a cooperative role for both T cell compartments in mediating tumor rejection **(Fig. 2f)**.

IFNγ ELISPOT analysis of TILs from mice treated with A20 neoVAX generated with mAars as the MHC-II neoAg and mLrrk1 revealed increased frequencies of mLrrk1- and mAars-specific T cells within the tumor compared with TILs isolated from tumor bearing mice treated with the irrelevant vaccine **(Fig. 2g)**. In addition, T cell responses against A20 antigens not included in the vaccine (i.e., mPrkcb and the A20 B-cell receptor) were increased exclusively in tumors derived from A20 neoVAX-treated mice **(Fig. 2g)**. These results suggest that effective neoAg vaccination enhances epitope spreading and broadens the antitumor T cell repertoire beyond the targeted epitopes.

Quantification using peptide-MHC tetramer staining further confirmed that combined MHC-I and MHC-II neoAg vaccination increased the absolute numbers of A20-specific CD4□ and CD8□ T cells relative to vaccination with either antigen alone **(Fig. 2h, i)**. Phenotypic analysis of mLrrk1-specific CD8□ T cells showed that inclusion of the A20 MHC-II neoantigen was associated with reduced expression of exhaustion markers of CD8^+^ T cells, as indicated by decreased frequencies of PD-1□CD39T cells, as and PD-1□ TIGIT□ cells **(Fig. 2j)**. This finding is consistent with synergistic enhancement of antigen-specific T cell responses when both MHC-I and MHC-II neoAg are included in the vaccine. Thus, engagement of both A20-specific CD4^+^ and CD8^+^ T cells induced by A20 neoVAX contributes to the maintenance of the functional state of tumor-specific CD8□ T cells during vaccine-induced tumor rejection.

### Combinatorial neoAg-based therapies are effective against immunotherapy-resistant subcutaneous A20 lymphoma

To determine whether checkpoint blockade could further augment vaccine-induced immunity, we evaluated the effects of the combination of A20 neoVAX with αPD-1 against subcutaneous A20 tumors **(Fig. 3a)**. Early αPD-1 treatment (days 6, 9, and 12 post-tumor inoculation) resulted in tumor rejection in approximately 20-26% of mice **(Fig. 1a)**, whereas delayed αPD-1 administration on day 15 failed to control tumor growth, confirming that established A20 subcutaneous tumors are largely insensitive to late PD-1 blockade **(Fig. 3b, c)**. The combination of an irrelevant vaccine administered on days 3, 10, and 17 together with delayed αPD-1 administration failed to induce tumor rejection. In contrast, combining the A20 neoantigen vaccine with αPD-1 significantly improved therapeutic efficacy, resulting in tumor rejection in 88.5% of tumor-bearing mice **(Fig. 3b, c)**. Thus, SLP neoantigen vaccines can sensitize otherwise resistant tumors to checkpoint therapy and the therapeutic benefit of such vaccines can be enhanced by combining them with ICT.

**Figure 3:**
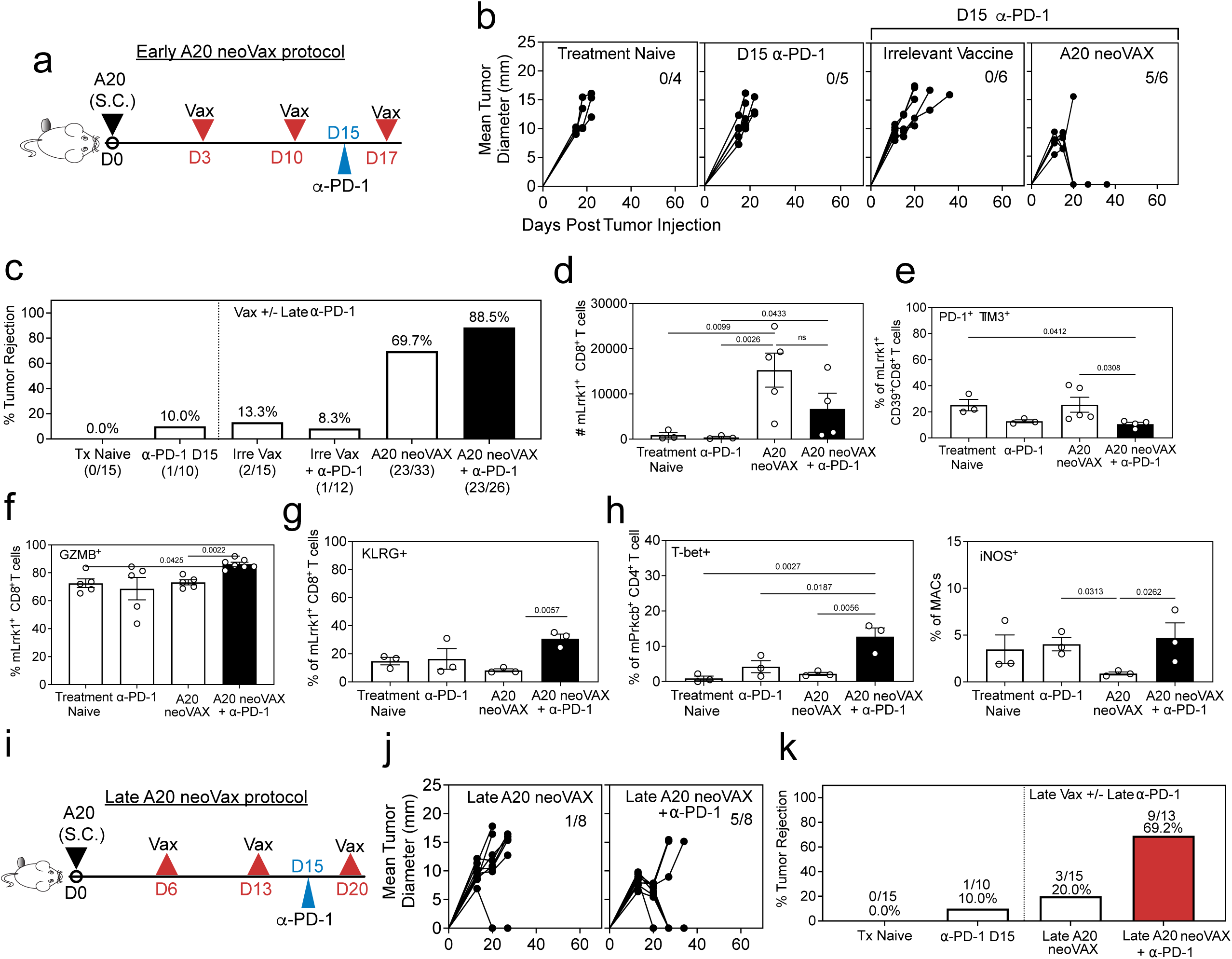
A20 neoVAX synergizes with late-PD-1 therapy to enhance subcutaneous A20 tumor control and T cell functionality. **a,** Schematic of the treatment protocol. **b,** Individual tumor growth curves of mice bearing subcutaneous A20 tumors treated as in panel **a**. Numbers indicate the number of mice in each group that rejected their tumors (complete responders) versus the total number of mice in each experimental group. **c,** Percent survival across treatment groups (combined from 5 independent experiments in A20 neoVAX or 3 independent experiments for vaccine plus αPD-1). **d,** Absolute numbers of mLrrk1-specific CD8□ T cells in tumors harvested on day 10 after tumor inoculation, **e,** Frequencies of mLrrk1-specific CD8□ T cells expressing CD39, PD-1 and TIM-3. **f-g,** Frequencies of mLrrk1-specific CD8□ T cells expressing granzyme B (**f),** and KLRG1**, (g)**. (**d-g**: Tx naïve / ⍰-PD-1, n=3; A20 neoVAX, n=5; A20 neoVAX + ⍰-PD-1, n=4). **h,** Frequencies of T-bet-expressing mPrkcb-specific CD4□ T cells **(right)** and iNOS-expressing macrophages **(left)** in tumors from indicated treatment groups (n=3 for each group).**i,** Experimental schematic of delayed neoantigen vaccination. **j-k,** Individual tumor growth curves **(j),** and summary of tumor rejection rates **(k),** following treatment as shown in i. (data are **combined** from 2 independent experiments). For **d-g**, data are shown as mean □±□ SEM, with each symbol representing one mouse. Statistical analyses were performed using one-way ANOVA with Tukey multiple-comparison correction.

The A20 neoVAX plus αPD-1 combination significantly increased the number of mLrrk1-specific CD8□ T cells within tumors compared with αPD-1 treatment alone **(Fig. 3d)**, suggesting enhanced expansion or recruitment of tumor-reactive CD8□ T cells. Relative to vaccine treatment alone, the combined treatment reduced the frequency of exhausted PD-1□ TIM3□ mLrrk1-specific CD8□ T cells **(Fig. 3e)** and increased effector CD8□ T cell populations, as reflected by higher frequencies of granzyme B- and KLRG1-expressing mLrrk1-specific CD8□ T cells **(Fig. 3f, g)**. These findings indicate that PD-1 blockade improves the functional quality of vaccine-induced CD8□ T cells. In addition to CD8□ T cell modulation, combined A20 neoVAX and αPD-1 increased the frequency of T-bet□ mPrkcb-specific CD4□ T cells and iNOS□ macrophages within tumors **(Fig. 3h)**, consistent with the induction of a Th1-skewed, inflammatory tumor microenvironment.

To assess the importance of vaccination timing, we next delayed the administration of A20 neoVAX to days 6, 13, and 20 post-tumor inoculation **(Fig. 3i)**. Delayed vaccination alone resulted in minimal tumor rejection, and late αPD-1 treatment alone remained ineffective. However, combining delayed neoantigen vaccination with late αPD-1 treatment significantly improved tumor control, leading to tumor rejection in 9/13 mice **(69.2%, Fig.3j, k)**. Although this efficacy was reduced compared with early vaccination plus αPD-1, delayed vaccination plus PD-1 achieved therapeutic efficacy comparable to early vaccination alone. These results indicate that PD-1 blockade can completely restore the effectiveness of vaccine-induced anti-tumor responses when vaccination is delayed and ineffective on its own.

Together, these data demonstrate that A20 neoVAX enhances both the magnitude and functional quality of tumor-specific immune responses, sensitizing immunotherapy-resistant A20 tumors to PD-1 blockade. The therapeutic efficacy of A20 neoVAX is reduced by delaying the timing of administration but can be restored by combination with ICT.

### The combination of neoantigen vaccination and ICT induces durable protection against systemic A20 lymphoma

To better recapitulate the pathology and progression of disseminated lymphoma, we established a systemic A20 lymphoma model and evaluated the therapeutic efficacy of neoantigen vaccination with or without ICT. To facilitate tumor detection, we used A20 cells expressing orange fluorescent protein (A20-OFP), enabling longitudinal in vivo and ex vivo monitoring. OFP expression did not confer detectable immunogenicity to A20, as A20 and A20-OFP cells exhibited comparable growth kinetics in immunocompetent mice in both subcutaneous and systemic settings **(Supplementary Fig. 3a,b)**.

In the systemic model, A20-OFP cells were injected intravenously and became detectable in the liver as early as 3 days post-injection **(Fig. 4a)**. Tumor cells then disseminated to the spleen and blood after day 17 **(Fig. 4a)**. Disease progression was associated with hepatosplenomegaly that was readily observed as a progressive increase in abdominal distension, increased body weight that was associated with tumor burden **(Fig. 4b-d and Supplementary Fig. 3c)**, and, at later stages (days 20-30), the development of severe morbidity. The distribution of tumor cells in the liver was not due to the tail vein injection route, as tumor cells were also observed in the liver following retroorbital injection **(Supplementary Fig. 3d)**.

**Figure 4:**
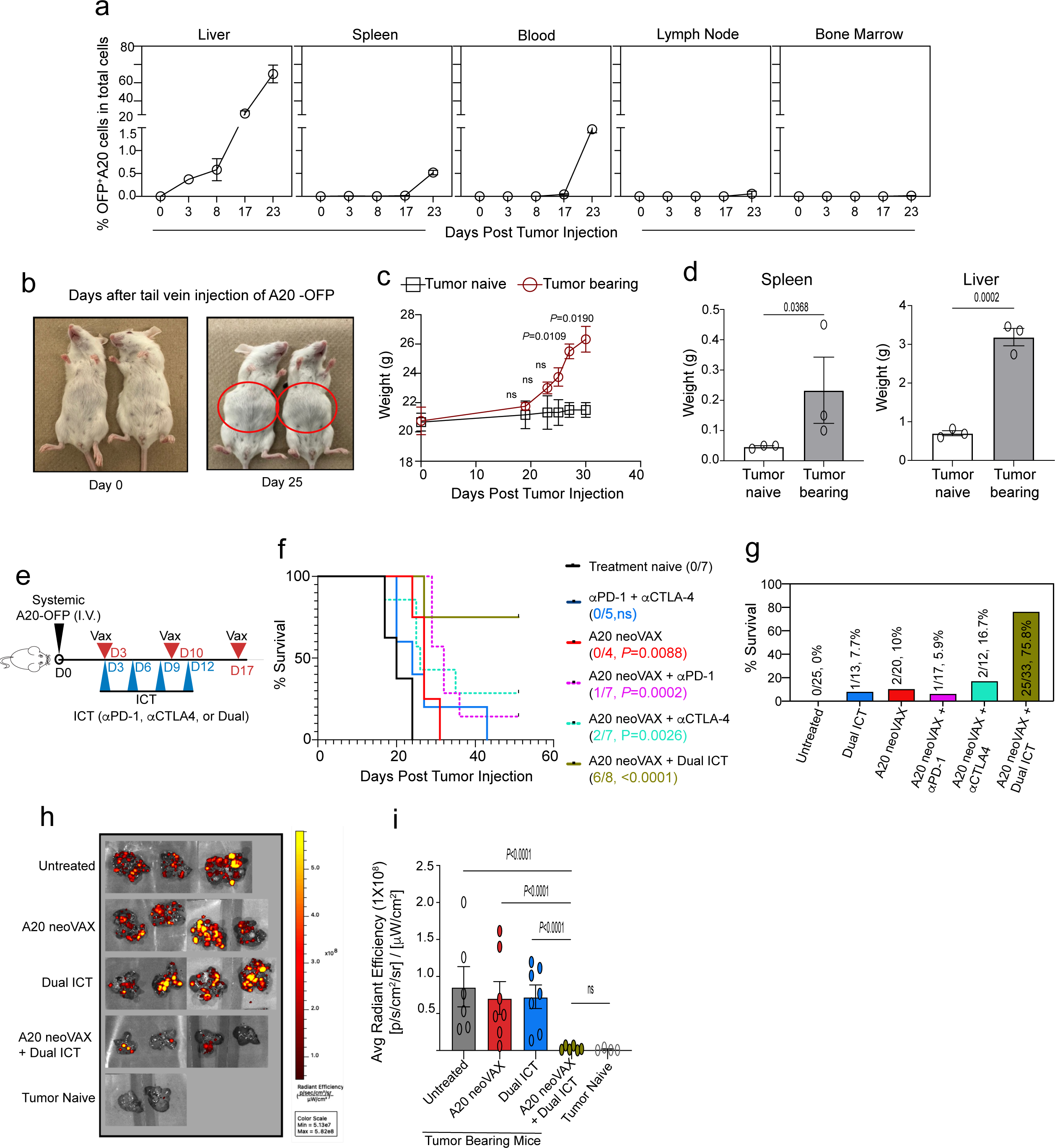
A20 neoVAX synergizes with dual ICT to eradicate systemic A20 lymphoma. **a,** Kinetics of A20-OFP tumor cell detection in liver, spleen, blood, lymph nodes, and bone marrow following intravenous tumor inoculation, quantified as the percentage of OFP□ A20 cells among total cells at indicated time points (n=3). **b,** Representative images of mice at day 0 and day 25 following intravenous injection of A20-OFP cells, illustrating the development of hepatomegaly caused by systemic tumor progression. **c,** Body weight over time in naïve and tumor-bearing mice following A20-OFP inoculation (tumor bearing, n=4; tumor naïve, n=6; two-way ANOVA Šídák’s multiple comparisons test). **d,** Weights of spleens (left) and liver (right) from naive and tumor-bearing mice at day 25 post tumor inoculation (n=3, Student’s t test). **e,** Experimental schematic for the treatment protocol for experiments through f-i. **f,** Representative Kaplan-Meier survival curves of mice receiving indicated treatments. Numbers of mice in each group and p values (calculated against the Tx-naïve group using the Mantel-Cox test) are shown. **g,** Percent survival of tumor-bearing mice across treatment groups (combined data from 5 independent experiments in Tx-naïve, A20 neoVAX + dual ICT; 2 experiments in A20 neoVAX + α-CTLA4, 3 experiments in all other groups). **h,** Representative ex vivo bioluminescence images of livers harvested on day 20 from tumor-bearing mice across treatment groups. **i,** Quantification of average radiant efficiency from ex vivo liver imaging in tumor-bearing and tumor-naive mice across treatment conditions (naïve, n=4; A20 neoVAX, n=7; others, n=6; one-way ANOVA with Tukey multiple comparison test). For **a, c, d,** and **i**, data are shown as mean □±□ SEM, with each symbol representing one mouse for **d** and **i**.

To assess therapeutic efficacy, mice bearing systemic tumors were assigned to six treatment groups and followed over time: (1) treatment-naive; (2) A20 neoVAX alone (MHC-I and MHC-II SLPs); (3) dual ICT (αPD-1□+αCTLA4); (4) A20 neoVAX □plus□αPD-1; (5) A20 neoVAX□plus αCTLA4; or (6) A20 neoVAX□plus□dual ICT. NeoAg vaccination was administered on days 3, 10, and 17 post-tumor injection, whereas single or dual ICT was administered on days 3, 6, 9, and 12 **(Fig. 4e)**.

Although, as presented above, dual ICT was sufficient to eradicate established *subcutaneous* A20 tumors, it provided no significant survival benefit in the *systemic* model **(Fig. 4f, g)**, indicating a fundamental difference in therapeutic sensitivity between localized and disseminated disease. A20 neoVAX alone (p=0.0088), as well as the vaccine combined with αPD-1 (p=0.0002) or CTLA4 (p=0.0026), significantly prolonged survival but did not prevent progressive tumor growth **(Fig. 4f, g)**. In contrast, combining A20 neoVAX with dual ICT resulted in a marked synergistic therapeutic efficacy, with approximately 75% of treated mice achieving durable tumor rejection (p=0.0001) **(Fig. 4f, g)**. The superiority of combinatorial therapy was further supported by ex vivo liver imaging and flow cytometric analyses, which revealed a substantial reduction in OFP^+^ A20 tumor cells in mice treated with A20 neoVAX plus dual ICT compared with all other treatment groups **(Fig. 4h, i)**. Importantly, none of the treatment regimens induced detectable adverse clinical effects in this model.

Collectively, these results demonstrate that neoantigen vaccination synergizes with combined PD-1 and CTLA4 blockade to induce durable and non-toxic rejection of ICT-resistant systemic lymphoma, establishing a rationale for neoantigen vaccine-based combinatorial immunotherapy in disseminated hematologic malignancies.

### Early A20 neoVAX intervention before immune checkpoint therapy is essential for effective combinatorial therapy against systemic A20 lymphoma

To further define the contribution of neoantigen vaccination to combinatorial immunotherapy, we first assessed whether both MHC-I and MHC-II neoantigens are required for therapeutic efficacy in the systemic A20 lymphoma model **(Fig. 5a)**. Combining an irrelevant neoantigen vaccine with dual ICT failed to induce systemic tumor rejection **(Fig. 5b, c)**. Vaccination with either the MHC-I or the MHC-II neoantigens alone, when combined with dual ICT, modestly delayed disease progression but did not prevent tumor outgrowth **(Fig. 5b, c)**. In contrast, the combination of both MHC-I and MHC-II neoantigens in the vaccine together with dual ICT resulted in robust therapeutic efficacy and durable survival, indicating that inclusion of both MHC-I and MHC-II neoantigens is required to achieve systemic A20 tumor rejection in the vaccine-ICT setting **(Fig. 5b, c).**

**Figure 5:**
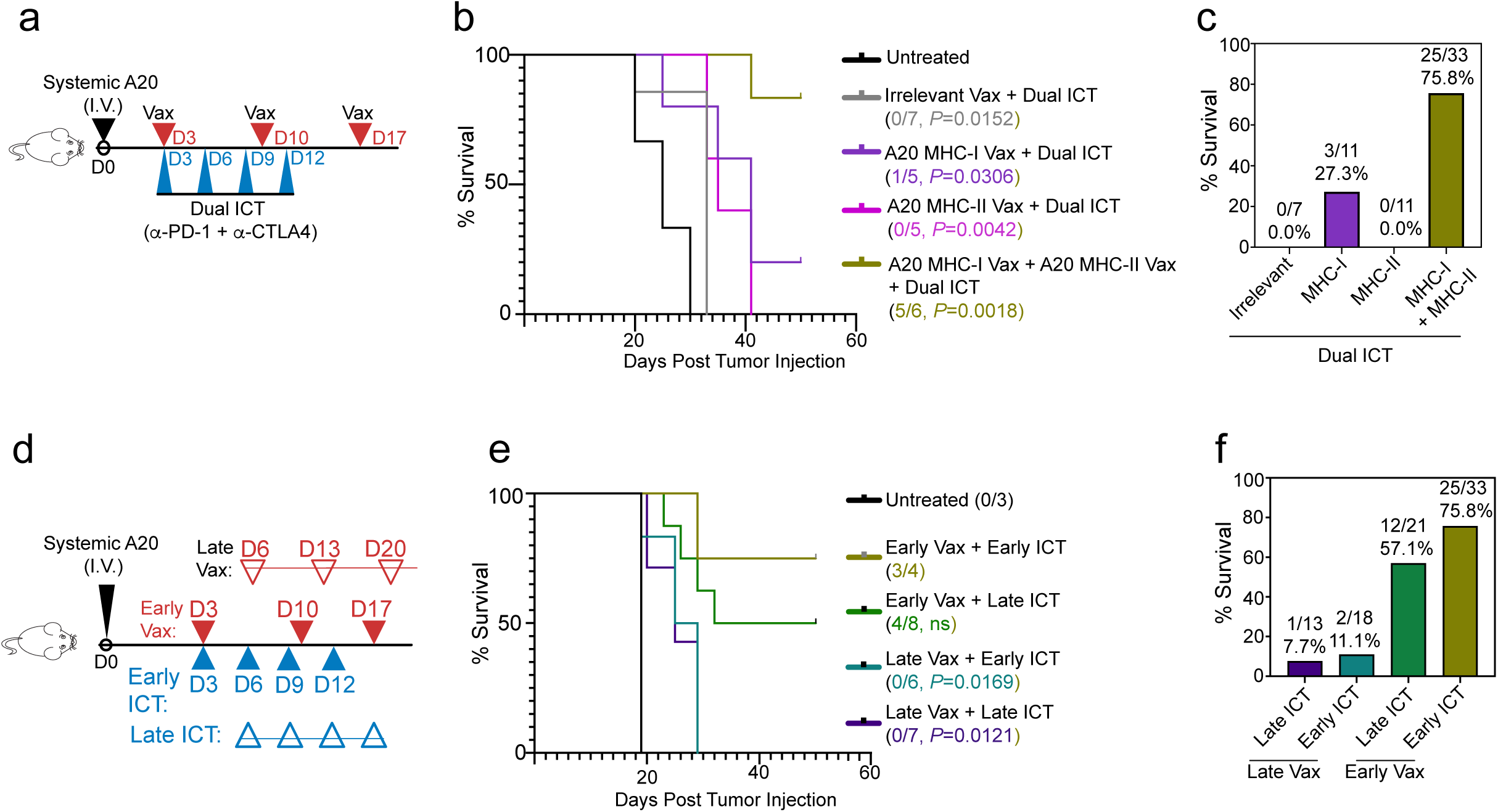
Early A20 neoVAX is required to achieve optimal synergy with dual immune checkpoint blockade in systemic A20 lymphoma. **a,** Experimental schematic. **b,** Kaplan-Meier survival curves of tumor-bearing mice treated as shown in **a** (*p* values are calculated against the Tx-naïve group using the Mantel-Cox test). **c,** Summary of survival outcomes across treatment groups treated as in **a**. **d,** Experimental schematic to study the impacts of treatment schedule on tumor outcome in **e-f**. **e,** Kaplan-Meier survival analysis (each individual condition is analyzed against the Early Vax + Early ICT group using the Mantel-Cox test). **f,** Percent survival across treatment groups stratified by neoAg vaccination and ICT timing (5 independent experiments for Early Vax + Early ICT group; 3 independent experiments for others).

To determine how treatment timing influences combinatorial efficacy, we next varied the temporal sequence of administering A20 neoVAX and dual ICT **(Fig. 5d)**. Early administration of A20 neoVAX (days 3, 10, and 17), followed by delayed dual ICT (days 6, 9, 12, and 15), effectively controlled systemic A20 lymphoma, resulting in approximately 60% survival of the treated group through day 50 **(Fig. 5e, f)**. In contrast, delaying neoantigen vaccination (days 6, 13, and 20) while administering ICT early (days 3, 6, 9, and 12) or late (days 6, 9, 12, and 15) failed to induce systemic tumor rejection **(Fig. 5e, f)**. Moreover, delaying both vaccination and ICT completely abrogated therapeutic efficacy, leading to progressive disease in all treated mice **(Fig. 5e, f)**.

Together, these results demonstrate that early neoantigen vaccination is a critical determinant of the success of combinatorial immunotherapy in systemic A20 lymphoma. While immune checkpoint therapy alone or when administered prior to vaccination is insufficient, early priming with neoantigen vaccines enables subsequent ICT to exert durable anti-tumor effects, highlighting the importance of treatment sequencing in neoantigen-based combinatorial strategies.

### A20 neoVAX and dual ICT synergize to prevent exhaustion and preserve the potency of A20 neoAg-specific CD8□ T cells locally and systemically

Whereas either combined immune checkpoint therapy or A20 neoVAX alone is sufficient to induce rejection of *subcutaneous* A20 tumors, effective clearance of *systemic* A20 lymphoma required the combination of A20 neoVAX and dual ICT. This distinction provided an opportunity to interrogate the mechanisms underlying vaccine-ICT synergy in the systemic setting.

Depletion of either CD4□ or CD8□ T cells abrogated the therapeutic efficacy of A20 neoVAX plus dual ICT, confirming that systemic tumor rejection also depends on both T cell subsets **(Supplemental Fig. 3e).** To characterize immune responses associated with therapeutic benefit, blood and liver samples were collected from treatment-naive mice and mice treated with A20 neoVAX alone, dual ICT alone, or the combination, and analyzed by flow cytometry at multiple time points **(Fig. 6a)**.

**Figure 6:**
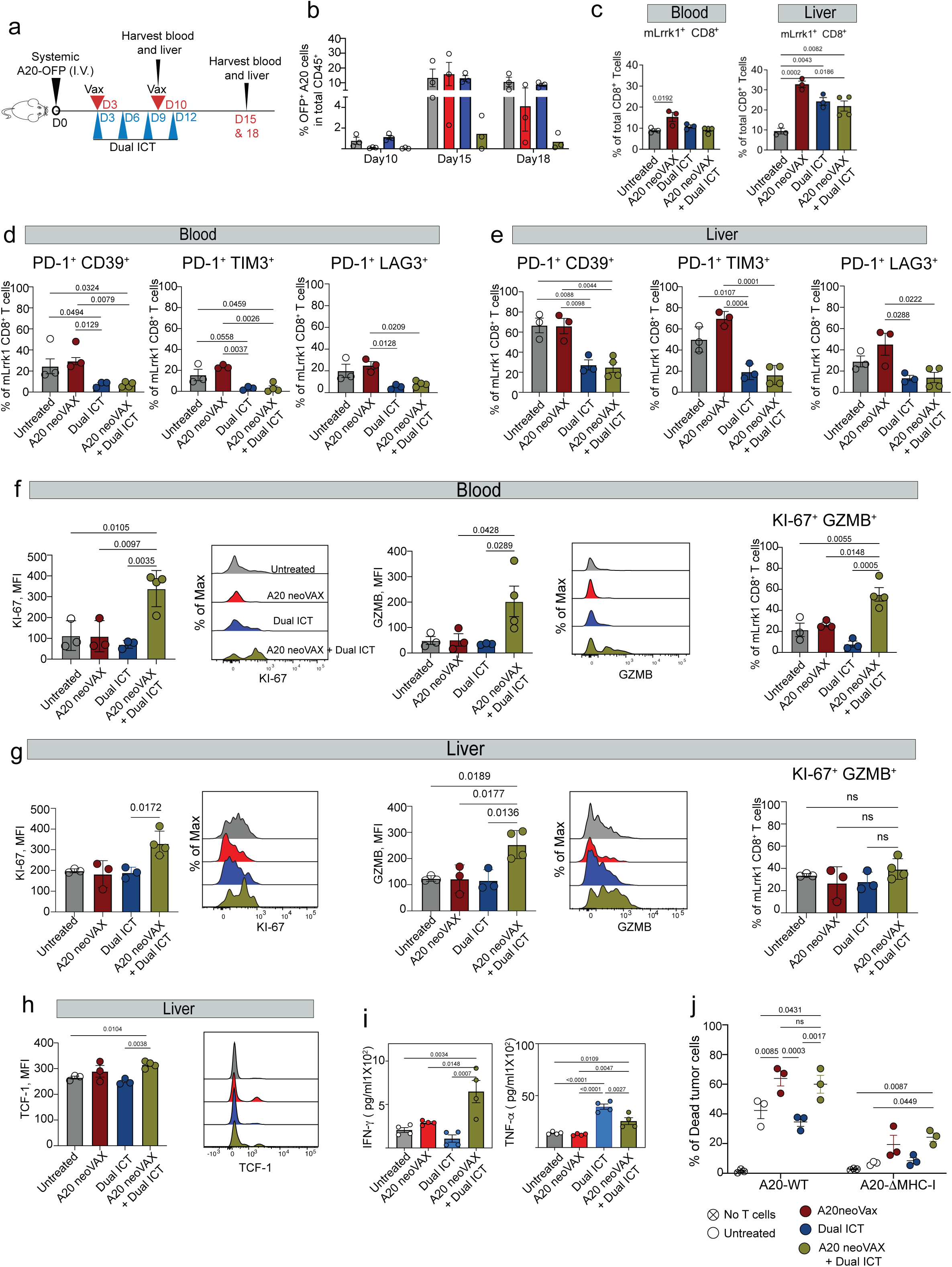
A20 neoVAX combined with dual ICT preserves functional tumor-specific CD8□ T cell responses and mediates systemic A20 tumor control. **a,** Experimental schematic indicating analysis of tumor-specific T cell responses in blood and liver following treatment with A20 neoVAX, dual ICT, or combination therapy. **b,** Frequency of OFP□ A20 tumor cells among total cells in the liver at the indicated time points following treatment. **c,** Frequencies of mLrrk1-specific CD8□ T cells among total CD8□ T cells in blood and liver across treatment groups 15 days post tumor injection. **d-h,** Phenotypic analysis of mLrrk1-specific CD8 T cells harvested on day 15 post tumor injections. **d, e,** Frequency of cells expressing PD-1 and CD39, PD-1 and TIM-3, or PD-1 and LAG-3 in blood **(d)** and liver **(e). f, g,** Expression of proliferation (Ki-67) and cytotoxicity (GZMB) markers in blood **(f)** and liver **(g)**. **h,** Expression of the stem-like marker TCF-1 in mLrrk1-specific CD8□ T cells in the liver. **i,** ELISA quantification for IFNγ and TNFα production by CD8□ TILs following ex vivo stimulation with 1μg/ml mLrrk1 peptide. **j,** Frequency of dead A20 tumor cells expressing NIR following co-culture with CD8□ TILs harvested from mice treated with multiple regimens (n=3 for each condition, two-way ANOVA with Tukey multiple comparisons test), comparing A20 WT and A20 ΔMHC-I targets. For **b-j**, data are presented as mean □±□ SEM; For **b-i**, each symbol represents one mouse, and statistical analyses were performed using one-way ANOVA with Tukey multiple-comparison correction.

We first examined tumor fate in livers at different time points across all four treatment regimens. At day 10 post-tumor inoculation, both A20 neoVAX alone and A20 neoVAX combined with dual ICT limited tumor expansion in the liver relative to treatment-naive and dual ICT-only groups **(Fig. 6b)**. By day 15, however, vaccine monotherapy no longer controlled tumor growth, whereas mice receiving combined A20 neoVAX and dual ICT exhibited markedly reduced tumor burden, ultimately leading to nearly complete tumor clearance **(Fig. 6b).**

At an early time point (day 10 post-tumor injection), although there were no significant differences in the frequency of mLrrk1-specific CD8^+^ T cells in the livers of treatment-naïve and treated mice (Supplementary Fig 4a), neoantigen vaccination significantly influenced the differentiation state of tumor-specific CD8□ T cells. Both A20 neoVAX alone and A20 neoVAX combined with dual ICT were associated with significantly lower frequencies of mLrrk1-specific CD8□ T cells expressing exhaustion-associated markers, including PD-1□CD39□, PD-1□TIM3□, and PD-1□LAG3□, compared with treatment-naive mice. PD-1 expression levels were also reduced in vaccination-containing groups **(Supplementary Fig. 4b).** A20 neoVAX alone significantly increased both the frequency and expression level of the stem-like transcription factor TCF1 in mLrrk1-specific CD8□ T cells compared with treatment-naive mice and A20 neoVAX plus dual ICT mice, consistent with induction of a progenitor-like CD8□ T cell state **(Supplementary Fig. 4c)**. In contrast, the addition of dual ICT to A20 neoVAX treatment prevented TCF1 upregulation and increased the frequency of CX3CR1□ and granzyme B-expressing mLrrk1-specific CD8□ T cells relative to vaccine alone, indicating an early skewing toward an effector-associated differentiation program **(Supplementary Fig. 4c).**

At a later time point (day 15), qualitative differences between treatment modalities became more pronounced. By day 15, A20 neoVAX, dual ICT, or the combination all increased the frequency of mLrrk1-specific CD8□ T cells in liver compared to untreated controls **(Fig. 6c)**. However, in both blood and liver, vaccine-alone-treated mice exhibited similar levels of exhausted PD-1□CD39□, PD-1□TIM3□, and PD-1□LAG3□CD8□ T cells comparable to treatment-naive mice, whereas dual ICT alone or combined with A20 neoVAX significantly decreased CD8□ T cell exhaustion **(Fig. 6d, 6e and Supplementary Fig. 4d)**. However, combined A20 neoVAX and dual ICT exclusively increased expression of granzyme B and Ki-67 on mLrrk1-specific CD8□ T cells in blood and liver, and a higher frequency of Ki-67□GZMB□mLrrk1-specific CD8□ T cells in blood **(Fig. 6f, g)**. Notably, combined treatment maintained elevated TCF1 expression in liver-infiltrating CD8□ T cells, beyond that observed in treatment-naive mice, suggesting sustained preservation of a stem-like CD8□ T cell pool **(Fig. 6h)**.

Functional assessment of CD8□ T cells further supported qualitative differences between treatment modalities. A20 neoVAX alone or combined with ICT increased the frequency of polyfunctional CD8□ T cells following short-term (4 hours) ex vivo antigen restimulation **(Supplementary Fig. 4e)**. However, mLrrk1-specific CD8□ T cells from mice receiving A20 neoVAX plus dual ICT produced significantly higher amounts of IFNγ and TNFα following prolonged (72□h) antigen restimulation compared with tumor-bearing mice treated with vaccine-alone **(Fig. 6i)**. In vitro cytotoxicity assays demonstrated enhanced MHC-I-dependent tumor cell killing by CD8□ T cells induced by A20 neoVAX either alone or in combination with dual ICT compared to CD8^+^ T cells harvested from untreated tumor-bearing mice or mice treated with dual ICT alone **(Fig. 6j)**. These findings indicate that neoantigen vaccination induces polyfunctional CD8□ T cell responses, while the addition of immune checkpoint therapy further enhances cytokine production without compromising cytotoxic activity.

Together, these results indicate that neoantigen vaccination and immune checkpoint blockade exert temporally distinct but complementary effects on tumor-specific CD8□ T cells. Early vaccination primes a stem-like CD8□ T cell population while limiting premature exhaustion, whereas subsequent immune checkpoint blockade redirects the fate of these vaccine-primed cells to prevent terminal exhaustion and sustain effector function. As a result, the combination of neoantigen vaccination and immune checkpoint therapy preserves both stem-like and effector CD8□ T cell compartments, enhances cytotoxic and cytokine-producing capacity, and enables durable systemic tumor control.

### A20 neoVAX and dual ICT synergize to enhance neoantigen-specific CD4□ T cell Th1 responses and mediate tumor rejection independent of tumor-intrinsic MHC-II expression

Given the indispensable role of CD4□ T cells in A20 neoVAX plus dual ICT-mediated systemic A20 tumor rejection, we next characterized neoantigen-specific CD4□ T cell responses across treatment groups. Significant differences were observed in the frequencies of mPrkcb-specific CD4□ T cells in the blood at day 10 but not at day 15 of mice treated with the combination of A20 neoVAX plus ICT. This combination consistently induced the highest frequencies of neoantigen-specific CD4□ TILs in the liver **(Fig. 7a and Supplementary Fig. 5a).**

**Figure 7:**
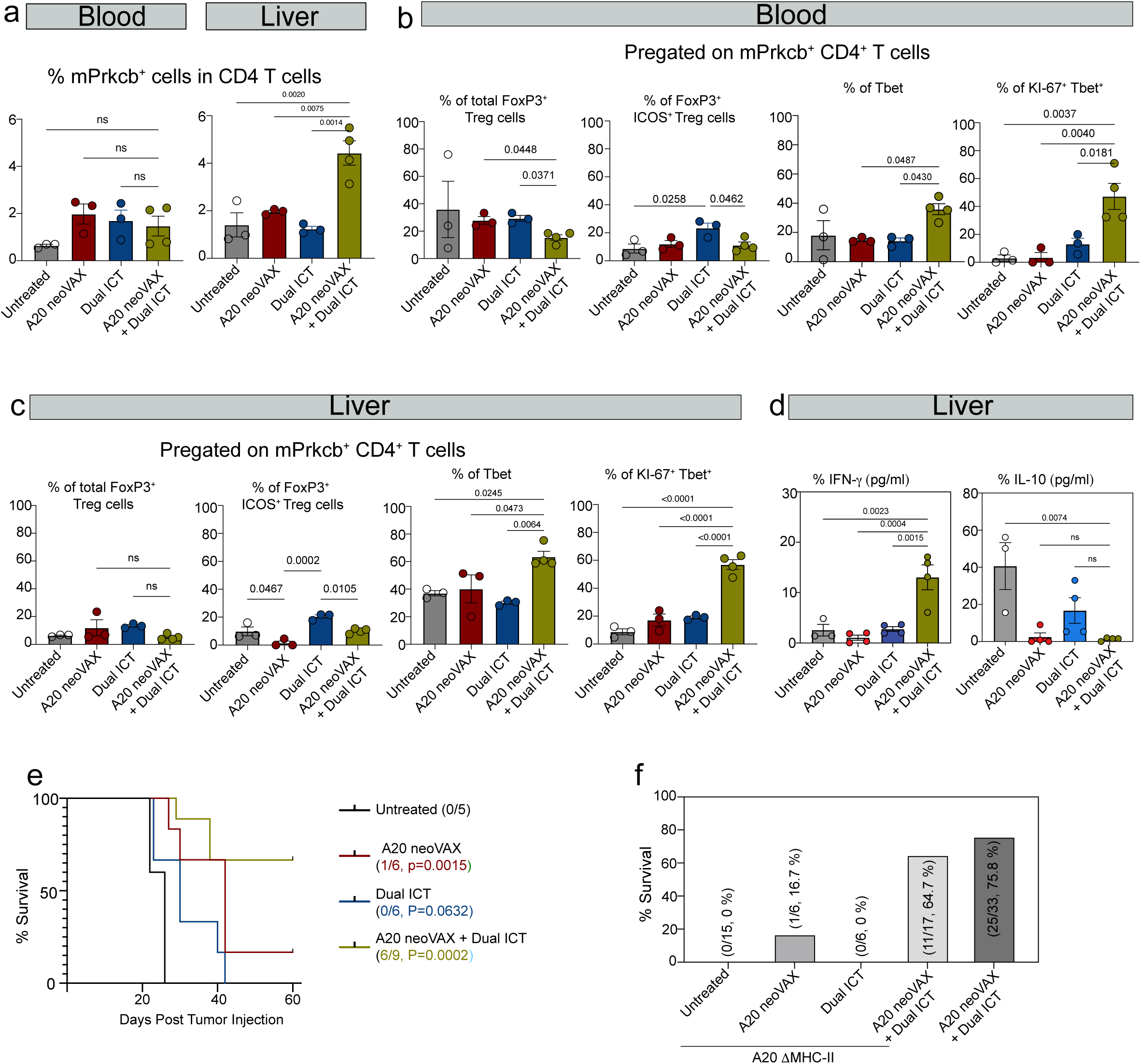
A20 neoVAX and dual ICT differentially shape neoantigen-specific CD4□ T cell responses in systemic A20 lymphoma. **a-d,** Blood and livers harvested from mice bearing systemic A20 tumors were harvested on day 15 post-tumor injection for detailed flow cytometric analysis of CD4 T cells. **a,** Frequency of mPrkcb-specific CD4□ T cells among total CD4□ T cells in blood and liver. **b-c,** Frequencies of total FOXP3□regulatory T cells (Tregs), FOXP3□ICOS□activated Tregs, T-bet□Th1 cells, and Ki-67□T-bet□ proliferating Th1 cells among mPrkcb-specific CD4□ T cells in blood **(b)** and liver **(c)** across treatment groups. **d,** ELISA quantification of IFNγ and IL-10 in supernatants following ex vivo restimulation of CD4□ TILs with 1μg/ml mPrkcb peptide for 72 hrs. **e-f,** WT BALB/c mice were intravenously injected with A20 ΔMHC-II lymphoma cells followed by different treatments. Kaplan-Meier survival analysis **(e)** as well as overall survival outcomes **(f)** are shown. For **a-d**, data are shown as mean □±□ SEM, with each symbol representing one mouse. Statistical significance was assessed using one-way ANOVA with Tukey multiple-comparison correction, with P values indicated in the panels. For **e**, statistical significance was assessed using log-rank (Mantel-Cox) tests for survival, with P values indicated in the panels.

Analysis of CD4□ T cell phenotypes revealed that total FoxP3^+^ regulatory T cell (Treg) frequencies were comparable across treatment groups in both blood and liver **(Fig. 7b, c)**. However, A20 neoVAX combined with dual ICT significantly limited the accumulation of ICOS□ Tregs compared with dual ICT alone. In contrast, combination therapy induced the highest proportion of T-bet□ Th1 CD4□ T cells in both blood (approximately 40% of mPrkcb-specific CD4□ T cells) and liver (approximately 60%) across all treatment groups **(Fig. 7b, c)**. Notably, in both blood and liver, more Ki-67□ Th1 cells were induced by A20 neoVAX plus dual ICT than by the other treatment, indicating synergistic expansion of proliferative Th1 CD4□ T cells with combination therapy **(Fig. 7b, c).** These findings suggest that A20 neoVAX and dual ICT cooperate to promote expansion of antigen-specific Th1 CD4□ T cells with enhanced proliferative capacity, a feature associated with effective CD8□ T cell support.

Functional profiling by intracellular cytokine staining further supported qualitative differences in CD4□ T cell responses. Following short-term (4 hours) ex vivo restimulation, CD4□ T cells from the A20 neoVAX plus dual ICT group displayed increased polyfunctionality and Th1 polarization, with higher frequencies of IFNγ□TNF□ cells compared with treatment-naive controls **(Supplementary Fig. 5b)**. After prolonged (72□h) antigen stimulation, supernatants were collected for cytokine measurement by ELISA. CD4□ T cells induced by A20 neoVAX plus dual ICT produced the highest levels of IFNγ across all treatment groups **(Fig. 7d)**. CD4□ T cells induced by A20 neoVAX alone and combination therapy produced low levels of IL-10 compared to CD4□ T cells from the untreated group **(Fig. 7d)**. Together, these data indicate that A20 neoVAX combined with dual ICT promotes a dominant Th1 cytokine program while limiting suppressive CD4□ T cell functions.

Loss of tumor-intrinsic MHC class II expression has been suggested to be an immune evasion mechanism in several human lymphomas and is associated with poor clinical outcomes^30,46–48^. A20 lymphoma cells intrinsically express high levels of MHC-II and are capable of antigen presentation, independent of IFNγ stimulation **(Supplementary Fig. 5d)**. To determine whether tumor-intrinsic MHC-II expression is required for the efficacy of A20 neoAg-based therapies, we generated MHC-II-deficient A20 cells and confirmed their deficiency in presenting mAars SLP to mAars T cell hybridoma **(Supplementary Fig. 5d, e)** and evaluated therapeutic responses in the systemic model. Similar to wild-type A20 tumors, systemic A20 tumors lacking MHC-II (A20ΔMHC-II) were efficiently rejected by the combination of A20 neoVAX and dual ICT, whereas neither treatment alone conferred long-term protection **(Fig. 7e-f)**. Alternatively, A20 neoVAX plus dual ICT increased the frequency of type 1 dendritic cells (cDC1) and enhanced MHC-II expression on these cells in systemic wild-type A20 tumors, suggesting that antigen presentation by professional antigen-presenting cells, rather than tumor cells, underlies effective CD4□ T cell priming in this model **(Supplementary Fig. 5f)**. We further confirmed tumor-intrinsic MHC-II independence in the subcutaneous A20 model, where both dual ICT alone and A20 neoVAX alone rejected A20ΔMHC-II tumors similarly to A20 wild type tumors **(Supplementary Fig. 5g, h)**.

Although CD4□ T cells expressing Granzyme B were detected in blood and liver and were enriched following A20 neoVAX plus ICT **(Supplementary Fig. 5c)**, in vitro cytotoxicity assays demonstrated that CD4□ T cells were unable to directly lyse A20 tumor cells, indicating that CD4□ T cells likely contribute to A20 tumor rejection through indirect mechanisms such as providing “help” to CD8^+^ T cells rather than direct cytotoxicity **(Supplementary Fig. 5i)**. Thus, taken together, these results establish that CD4□ T cells play a critical role in the efficacy of neoAg-based therapy for A20 lymphoma by providing help to tumor-specific CD8^+^ T cell development and function.

### CD8-targeted cytokines improve the efficacy of A20 neoVAX

A20 neoVAX plus checkpoint blockade increased CD4-derived production of effector cytokines while reducing IL-10 in systemic A20 lymphoma. These findings thus suggest that productive systemic A20 control may be associated with a specific cytokine environment that reinforces CD8 T-cell immunity. We next asked whether direct cytokine support of the CD8^+^ T-cell compartment could further improve therapeutic efficacy. More importantly, given the potential toxicity of □CTLA4 in clinical practice,^49^ other therapies that can efficiently induce rejection of systemic A20 tumors may be more favorable for clinical development as a substitute for □CTLA4.

IL-2 has long been recognized as a crucial cytokine for T cell proliferation, function, and maintenance^50^. Indeed, A20 neoVAX and dual ICT synergized to induce the highest amount of IL-2 secretion from A20-specific CD4^+^ T cells compared to single-agent therapies **(Fig. 8a)**. However, the systemic use of IL-2 is limited by its broad and sometimes toxicity-promoting activity on multiple immune subsets, including regulatory T cells, innate lymphoid populations, and NK cells^51–53^. Recent work has described a murine CD8-targeted IL-2 surrogate (CD8-IL2 or AB248), a cis-targeted fusion protein comprising a CD8-targeting antibody and an IL-2 mutein with ablated IL-2Rα binding and attenuated IL-2Rγ binding. This design enables preferential IL-2 signaling on CD8^+^ T cells, promoting their expansion, effector differentiation, cytotoxicity, and antitumor activity while limiting activation of Tregs, NK cells, and other non-CD8 IL-2R-expressing populations^44,54,55^. The human version of this reagent has been shown to promote clinical benefit as a monotherapy in human cancer clinical trials with limited major toxicities^56^.

**Figure 8:**
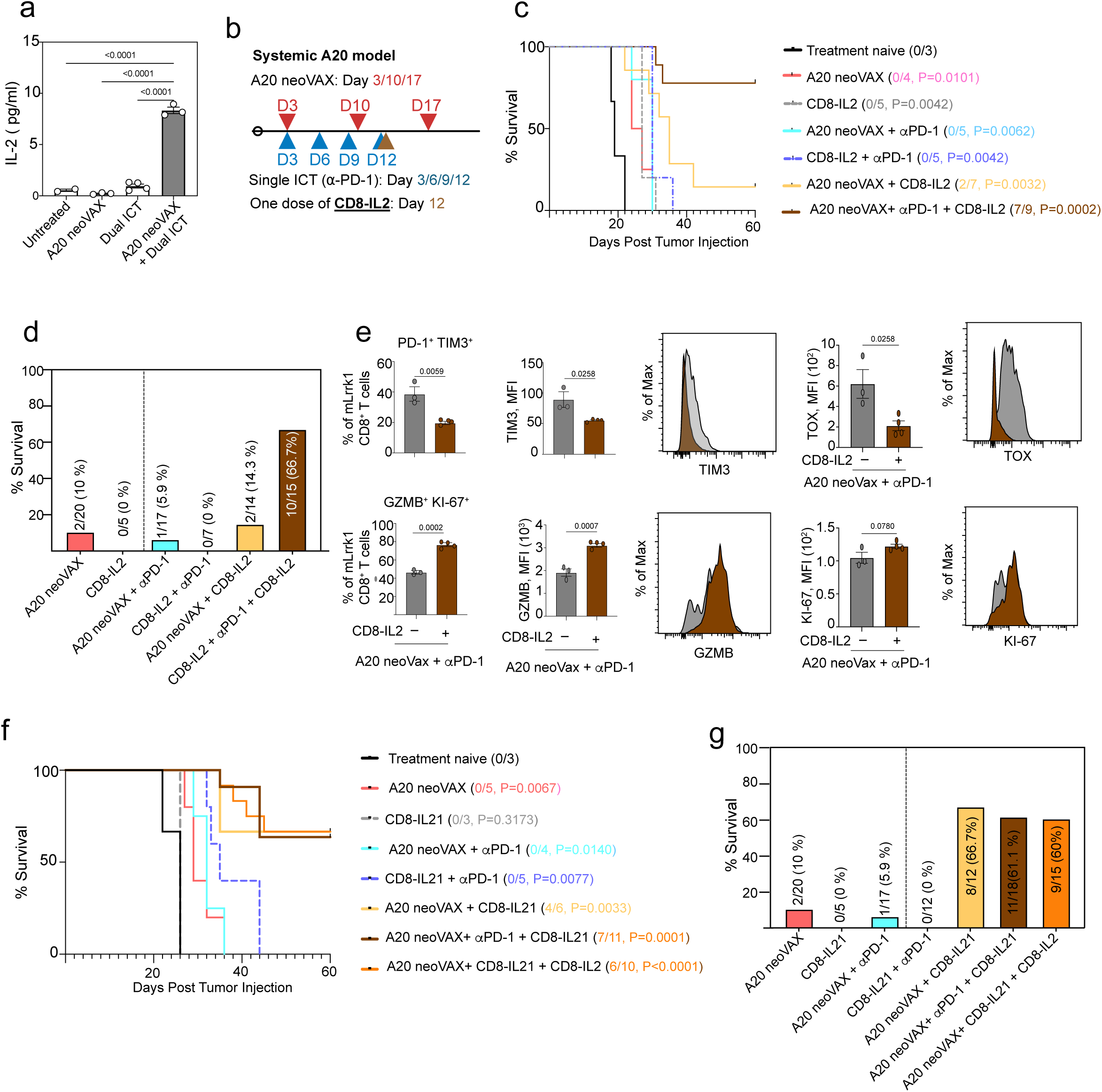
CD8-targeted IL2 (CD8-IL2) and CD8-IL21 enhance neoantigen vaccine-based therapy in systemic A20 lymphoma. **a,** ELISA quantification of IL-2 in supernatants following ex vivo restimulation of CD4□ TILs with 1μg/ml mPrkcb peptide for 72 hrs. **b,** Treatment schedule. **c,** Kaplan-Meier survival curves of tumor-bearing mice treated as shown in b (*p* values are calculated against the Tx-naïve group using the Mantel-Cox test). **d,** Summary of survival outcomes across treatment groups. **e,** Flow cytometry analysis of mLrrk1-specific CD8□ T cells from mice treated with neoVAX plus α-PD-1 in the presence or absence of CD8-IL2. Frequencies of PD-1□ TIM3□ and GZMB□ KI67□ cells and mean fluorescence intensity (MFI) of TIM3, TOX, GZMB, and Ki-67 are shown, together with representative histograms. **f,** Tumor-bearing mice were treated as in **b** but with CD8-IL21 and Kaplan-Meier survival analysis **(f)** and pooled survival frequencies across the treatment groups (**g)**. (*P* values were calculated by comparing each group with the Tx-naïve group using the Mantel–Cox (log-rank) test.).

We first tested whether combining CD8-IL2 with A20 neoVAX could effectively control systemic A20 tumors. Mice bearing systemic A20 lymphoma were treated with A20 neoVAX on days 3, 10, and 17, αPD-1 on days 3, 6, 9, and 12, and a single dose of CD8-IL2 on day 12 **(Fig. 8b)**. CD8-IL-2 alone, or in combination with αPD-1, or together with A20 neoVAX failed to protect mice from systemic A20 lymphoma. In contrast, the combination of A20 neoVAX + PD-1, + CD8-IL2 substantially improved therapeutic efficacy, resulting in long-term survival in 66.7% of mice **(Fig. 8c, d)**. These data indicate that whereas CD8-IL2 signaling is not sufficient as a monotherapy in systemic A20 lymphoma, it nevertheless cooperates with neoantigen vaccination and PD-1 blockade to promote durable tumor control.

Immune profiling showed that the addition of CD8-IL2 therapy improved the phenotype of vaccine-primed antigen-specific CD8 T cells. In mice treated with A20 neoVAX + αPD-1, adding CD8-IL2 into the treatment reduced exhaustion-associated features, including decreased PD-1□ TIM3□ cells and reduced TIM3 and TOX expression **(Fig. 8e)**. At the same time, CD8-IL2 therapy enhanced effector-associated features, including increased GZMB expression and a higher frequency of GZMB□ Ki67□ mLrrk1-specific CD8 T cells **(Fig. 8e)**. These data indicate that CD8-IL2 does not merely expand CD8 T cells, but improves the functional state of the vaccine-induced tumor-specific CD8 T cell pool.

Recently, IL-21 has been shown to support CD8 T cell memory formation, sustain function during chronic antigen exposure, and improve the efficacy of PD-1 or CTLA4 blockade in preclinical tumor models^57–59^. More recent work further demonstrated that CTLA4-based checkpoint therapy depends on IL-21 signaling to mediate cytotoxic reprogramming of PD-1□ CD8□ T cells, supporting IL-21 as a rational cytokine-based strategy for augmenting checkpoint-responsive T cell states^60^. Based on these observations, we tested a murine CD8-targeted IL-21 surrogate (CD8-IL21 or AB821) in the systemic A20 model. CD8-IL21 is a cis-targeted fusion protein comprised of a mouse CD8-targeting antibody and an IL-21 mutein with attenuated IL-21R binding and a reduced positive-charge profile. This design improves bioavailability and enables avidity-dependent IL-21 activity preferentially on CD8^+^ T cells while limiting activity on non-CD8^+^ IL-21R-expressing cells^61–63^. Similar to CD8-IL2, CD8-IL21 alone or when combined with αPD-1 failed to produce long-term survival. However, combining A20 neoVAX with CD8-IL21 markedly improved survival, with 66.7% of mice rejecting their tumors and surviving long term **(Fig. 8f, g)**. Impressively, these results did not require the addition of αPD-1. Thus, CD8-IL21 potentiates the efficacy of the A20 neoantigen vaccine, even in the absence of PD-1 blockade, thus supporting the broader conclusion that effective systemic lymphoma control requires both antigen-specific priming and cytokine-mediated reinforcement of CD8 T cell function. Of note, the addition of CD8-IL2 to the A20 neoVAX + CD8-IL21 treatment did not increase therapeutic efficacy **(Fig. 8f, g)**, suggesting that the two CD8-directed muteins functioned through overlapping but not identical mechanisms.

Together, these findings extend the therapeutic potential of personalized cancer vaccines. Systemic A20 lymphoma cannot be effectively controlled by checkpoint blockade, neoantigen vaccination, or cytokine treatment in isolation. Durable rejection requires coordinated immunotherapy in which A20 neoVAX establishes tumor-specific T cell immunity, αPD-1 protects the response from dysfunction, and CD8-targeted cytokines reinforce CD8^+^ T cell effector fitness. Thus, CD8-targeted cytokines, and CD8-IL21 in particular, because it enhances vaccine efficacy without concomitant PD-1 blockade in this model, represent rational strategies for improving neoantigen vaccine efficacy in the systemic A20 B-cell lymphoma model. These targeted cytokines safely convert vaccine-primed CD8^+^ T cells into more functional, less exhausted anti-tumor effectors, potentially even without needing immune checkpoint therapy. The translational potential of these targeted cytokines is supported by published phase 1 safety and tolerability data for the CD8-IL2 clinical program^56^ and by preliminary tolerability findings from the ongoing phase 1 evaluation of the CD8-IL21 clinical program^61^.

## DISCUSSION

Personalized neoantigen vaccines represent a novel approach to generate tumor-specific T cell responses and induce favorable clinical outcomes in solid tumors, but their therapeutic potential in B-lymphoma remains poorly defined^1–8^. Early therapeutic lymphoma vaccines have largely focused on lymphoma cell idiotypes, targeting the variable regions of their B-cell receptors^64–66^. Both preclinical studies and clinical trials have demonstrated the safety and feasibility of idiotypic lymphoma vaccines with meaningful immune correlations^64–72^. With advances in vaccine platforms (antigen selection, vaccine formats, and adjuvants) and increased understanding of tumor neoantigens over the past decades, the question of whether targeting lymphoma neoantigens with vaccine-based approaches could be an effective therapeutic strategy has thus become an increasingly important topic for advancing lymphoma therapy design^5,8–11^. A recent pilot clinical study has demonstrated that personalized neoantigen vaccines can be a safe and feasible approach and presented promising clinical outcomes when combined with αPD-1 in patients with refractory B cell lymphoma^10^. However, the optimal design of neoantigen vaccines (requirements for antigen composition, efficacy of monotherapy vs. combinatorial therapy, possible therapeutic combinations, etc.) required further evaluation.

Here, using syngeneic subcutaneous and systemic A20 B-cell lymphoma models, we show that A20 neoVAX can mediate tumor control as monotherapy in localized disease and can cooperate with immune checkpoint therapies and targeted cytokine therapies to produce durable rejection of orthotopically-administered disseminated lymphoma. Our findings identify several determinants of efficacy of neoAg-based therapies in both the subcutaneous and systemic lymphoma models, including coordinated targeting of MHC-I and MHC-II neoAg, synergy between neoAg vaccines and other immunotherapies (treatment timing and order), maintenance of functional lymphoma-specific CD4□ and CD8□ T cell responses by neoAg-based therapies, and introduction of CD8-targeted cytokines as a crucial combination with lymphoma therapeutic vaccines.

Despite the widespread use of the A20 tumor cell line to model lymphoma therapy, most publications have utilized subcutaneous A20 tumor localization^12–17^. In the current study, we established both subcutaneous and systemic A20 models to better recapitulate disease in a more physiologic setting and thereby examine the therapeutic efficacy of A20 neoVAX. It is now well established that tumor cell lines display distinct sensitivities to immunotherapy when injected into naïve mice either orthotopically or at non-physiologic sites^73,74^. We observed a similar divergence in the A20 model, highlighting the importance of evaluating immunotherapies in orthotopic settings that more closely reflect the immunological context of tumor development and progression. These differences nonetheless demonstrate the therapeutic activity of A20 neoVAX in both localized and systemic disease settings, despite the additional barriers to immune control imposed by disseminated tumor distribution and distinct tissue immune environments.

Mechanistically, effective lymphoma neoAg-based vaccine therapy requires the presence of both MHC-I and MHC-II neoAgs. Whereas vaccines employing only MHC-I or MHC-II antigens by themselves did not support durable tumor rejection (in either subcutaneous or systemic A20 tumor models), vaccines comprised of the combination of MHC-I and MHC-II neoantigens were efficacious. These findings support a model in which MHC-I neoAgs promote CD8□ T cell effector responses, while MHC-II neoAgs support CD4□ T cell functions required for sustained anti-tumor immunity, likely mediated through helper functions that support antigen presentation, CD8□ T cell maintenance, and remodeling of the tumor microenvironment^22,24,45^. Importantly, A20-intrinsic MHC-II expression is not required for the efficacy of A20 neoVAX-based therapies in both subcutaneous and systemic A20 models. A20ΔMHC-II tumors remain as susceptible to A20 neoVAX-based therapies as WT A20 tumors, suggesting that MHC-II-restricted CD4□ T cell responses can be supported through host antigen-presenting cells. This finding may be relevant to human lymphomas, despite the fact that MHC-II loss is frequently associated with immune evasion and poor clinical outcomes^30,46–48^.

Several effective therapeutic combinations have been examined here. While ICT has achieved limited success in NHL^33–35^, we demonstrate that a combination of A20 neoVAX and ICT can synergistically provide durable immune protection against established A20 tumors, consistent with the observations from several key clinical neoantigen vaccine studies in solid tumors^1–4,24–26^. Importantly, the therapeutic efficacy of A20 neoVAX plus ICT can be achieved comparably by combining A20 neoVAX with the two novel CD8-targeting cytokines, CD8-IL2 or CD8-IL21, where either agent was able to prevent exhaustion and sustain proliferative and cytotoxic CD8^+^ T cell states. Of note, we observe differences in the requirement for αPD-1 for the therapeutic effectiveness of A20 neoVAX when combined with CD8-IL2 versus CD8-IL21. Whereas A20 neoVAX plus CD8-IL2 is effective only when used with αPD-1 in systemic therapy models, A20 neoVAX plus CD8-IL21 can clear systemic A20 tumors without αPD-1. This difference may be partially explained by the distinct downstream signaling pathways of CD8-IL2 and CD8-IL21. Although the receptors for both cytokines share the common γ chain (γc) and activate Jak1 and Jak3, IL-2 preferentially activates Stat5, whereas IL-21 preferentially and more durably activates STAT3, together with context-dependent activation of STAT1 and STAT5^50,75^. A previous report has proposed that IL-2 and IL-21 promote distinct differentiation programs in CD8^+^ T cells based on studies that employed adoptive cell therapy models^76^. Whether the same is true for direct in vivo targeting to CD8^+^ T cells remains to be seen. Our experiment showing no synergy or functional interference between the addition of CD8-IL2 plus CD8-IL21 to A20 neoVAX would rather suggest that these two related cytokines promote overlapping but nonidentical functions to tumor-specific CD8 T cells. More work will be needed to fully define the common versus distinct mechanisms involved in the action of these two important cytokines.

While non-targeted IL-2 and IL-21 have limited use as systemic therapies due to their pro-inflammatory and immunosuppressive effects on non-CD8^+^ T cells, our preclinical findings in lymphoma models demonstrate that either CD8-IL2 or CD8-IL21 safely achieves superior therapeutic efficacy when combined with lymphoma neoAg vaccines. Our study also reveals that B-lymphoma cells, which were thought to be non-immunogenic due to constant cancer immunoediting because of their residency in immune compartments, can indeed harbor sufficient mutational neoantigens so as to become targets of adaptive immunity, leading to their therapeutic elimination under proper circumstances. Taken together, the results of this study provide strong experimental support for further translational efforts in human B-lymphoma.

Overall, our findings establish that neoAg vaccination can generate coordinated CD4□ and CD8□ T-cell immunity in B-lymphoma and that rational combinations with other immunotherapies can convert this response into durable systemic control of B-lymphoma. More importantly, while most previous neoAg vaccine studies in other cancer types have focused on combining neoAg vaccines with traditional immunotherapies such as ICT^1–4,24–26^, our results bring novel targeted cytokine therapies into focus and show that they achieve anti-tumor efficacy comparable to ICT when either approach is combined with neoAg vaccines. Translationally, these results provide a framework for optimizing previously piloted neoAg vaccine-based immunotherapy in B-lymphoma^10^ and identify mechanistic determinants that may inform the design of clinically translatable combination strategies.

## Figures and Figure Legends

**Supplementary Figure 1:**
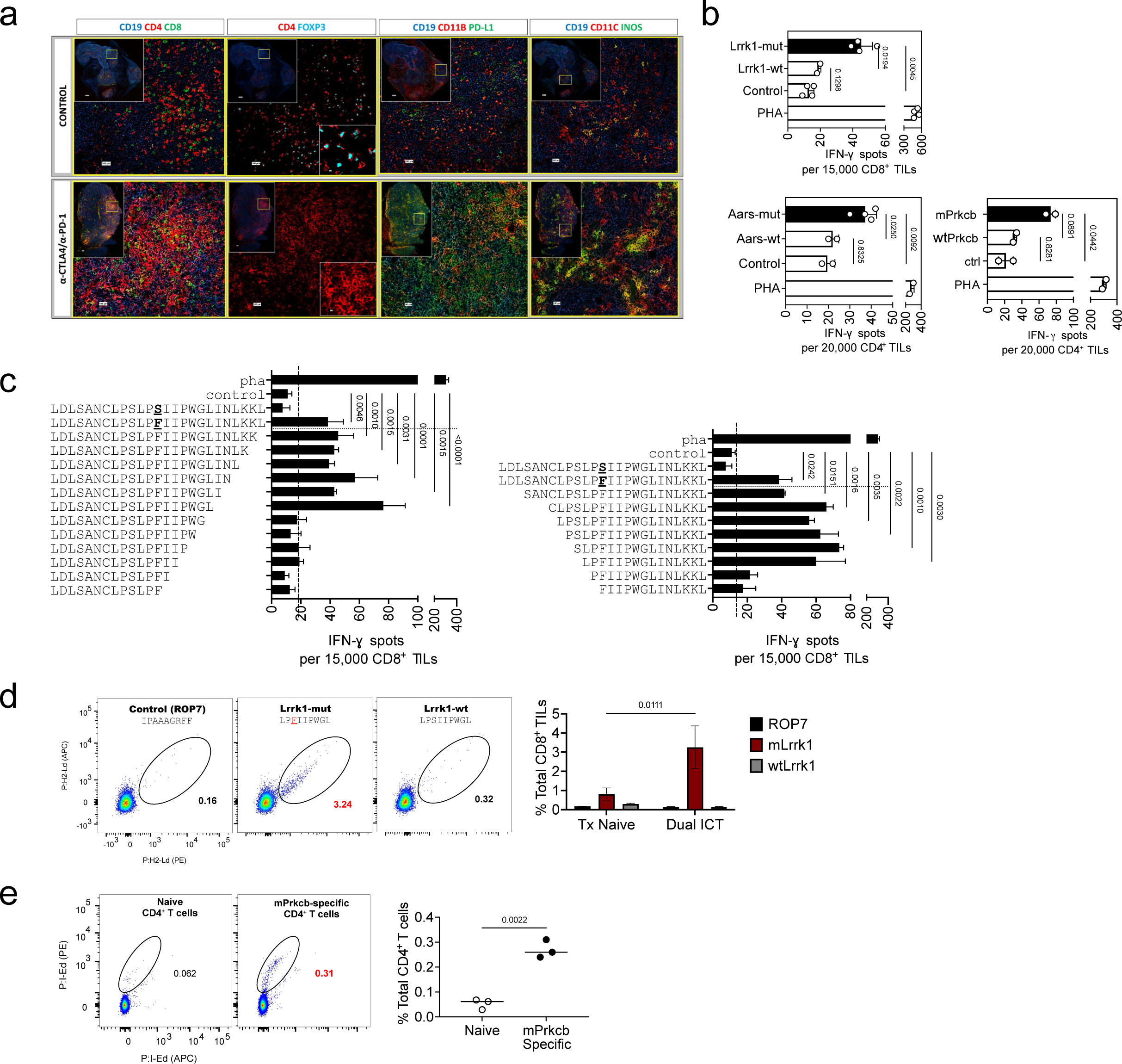
Validation of T cell responses specific to A20-expressed neoantigens. **a,** CODEX multiplex imaging showing the spatial reorganization of selected immune cells in subcutaneous A20 lymphoma tumors after α-CTLA-4/α-PD-1 immunotherapy. BALB/c mice bearing A20 tumors were treated with either Control Ab or α-CTLA-4/α-PD-1, and tumors were harvested 13 days later, fresh frozen, cut into 8 γm thick tissue slices, and stained with our expanded CODEX antibody panel. CD19 is used as a marker for A20 lymphoma cells. **b,** IFN ELISpot analysis of tumor-infiltrating CD8□ /CD4□ T cells harvested from dual ICT-treated tumor-bearing mice following stimulation with mutant A20 predicted peptides or their corresponding wild-type sequences. Data are shown as mean□±□ s.e.m. from n = 2-4 biological replicates (one-way ANOVA with Tukey’s multiple comparisons test). **c,** IFNγ ELISpot of tumor-infiltrating CD8□ T cells isolated from dual ICT-treated tumor-bearing mice following stimulation with truncated mutated Lrrk1 peptides and control wild-type Lrrk1 peptide (n=2). **d,** Representative flow cytometry plots **(left)** and corresponding quantification (**right**) of CD8 TILs stained with different peptide-MHC-I monomers labeled with PE-streptavidin or APC-streptavidin. (right, n=2, two-way ANOVA with Šídák’s multiple comparisons test). **e,** Representative flow cytometry plots **(left)** and corresponding quantification (**right**) of CD4 TILs stained with mPrkcb-MHC-II monomers labeled with PE-streptavidin or APC-streptavidin (n=3, one-way ANOVA with Tukey’s multiple comparisons test).

**Supplementary Figure 2:**
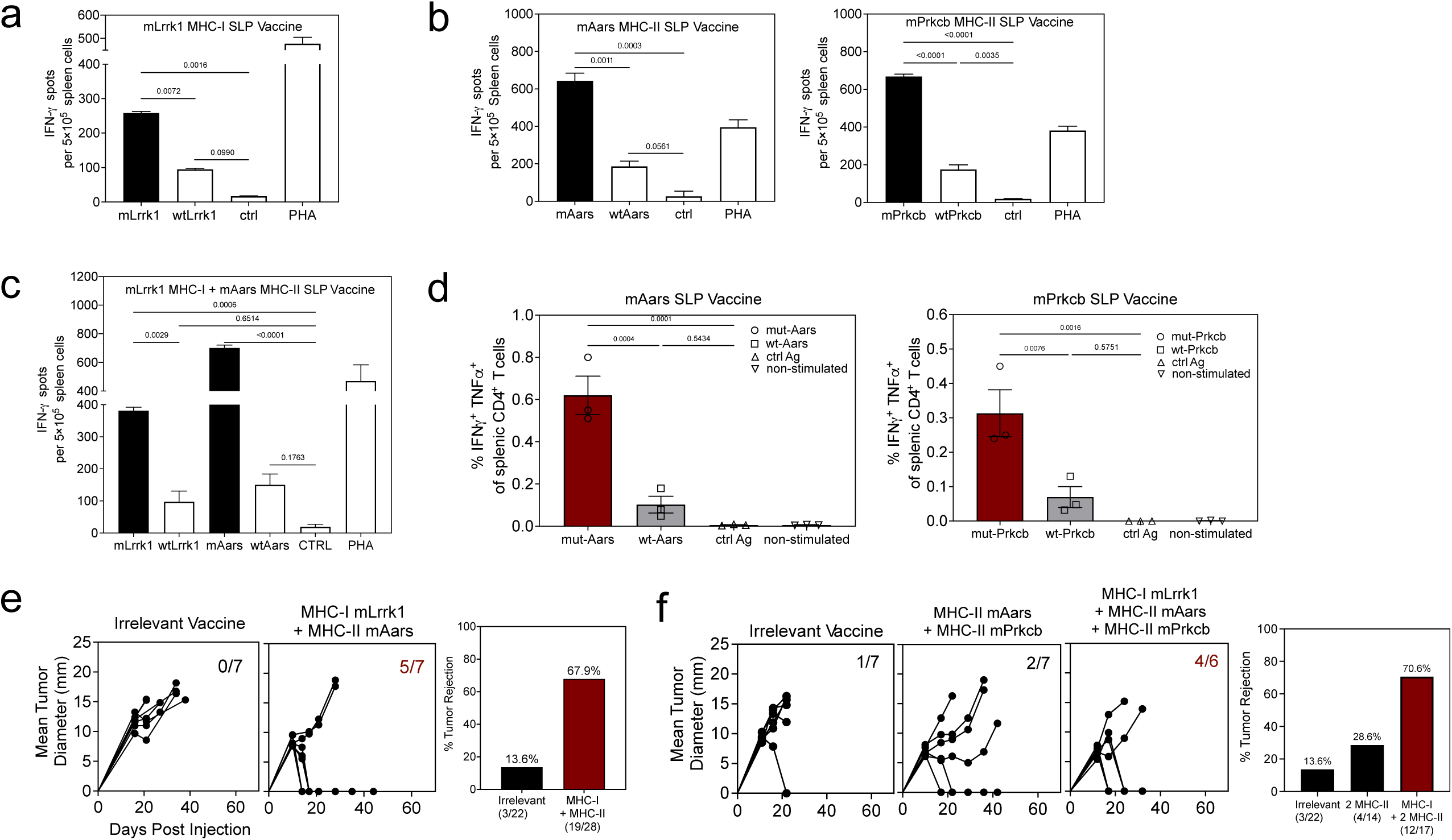
SLP vaccines encoding A20 neoantigens induced mutation-specific responses in vivo. **a,** IFNγ ELISpot analysis of splenic cells from mice treated with A20 MHC-I mLrrk1 SLP + poly-ICLC following stimulation with 10 γM of the indicated mutant or corresponding wild-type peptides. PHA was used as a positive control at 1 γg/ml (n=2, one-way ANOVA with Tukey’s multiple comparisons test). **b**, IFNγ ELISpot analysis of splenic cells from mice treated with A20 MHC-II mAars or mPrkcb SLP + poly-ICLC following ex vivo stimulation with 10 γM of the indicated mutant or corresponding wild-type peptides (n=2, one-way ANOVA with Tukey’s multiple comparisons test). **c,** IFNγ ELISpot analysis of splenic cells from mice treated with vaccines containing both MHC-I and MHC-II neoAg vaccines following ex vivo stimulation with 10 γM of the indicated mutant or corresponding wild-type peptides (n=2, one-way ANOVA with Tukey’s multiple comparisons test). **d,** Frequency of IFNγ and TNFα expressing CD4 T cells isolated from mAars and mPrkcb SLP-vaccinated mice following ex vivo stimulation with the indicated peptides for 4 h in the presence of Golgi Plug, followed by intracellular staining (n=3, one-way ANOVA with Tukey’s multiple comparisons test). All data are shown as mean □±□ s.e.m. **e,f.** Representative tumor growth curves **(left)** and pooled percent survival **(right)** of tumor-bearing mice treated with the indicated vaccines. The number of mice that rejected their tumors relative to the total number of mice in each group is indicated (Data are representative of 4 independent experiments for the MHC-I+MHC-II group and 3 independent experiments for the irrelevant vaccine group).

**Supplementary Figure 3:**
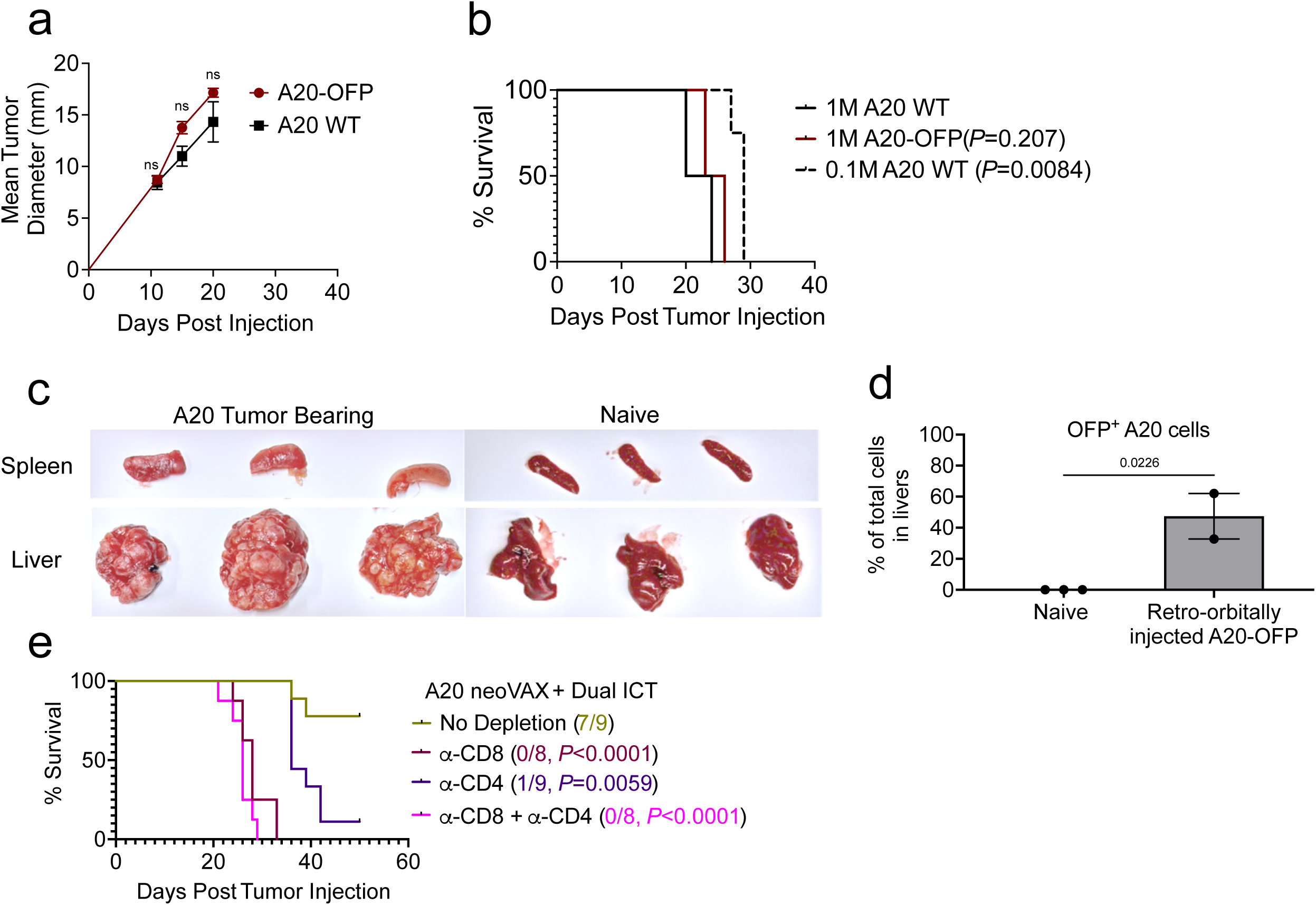
Comparison of systemic A20 model established by different cell lines and injection routes. **a,** Tumor growth curves of A20 WT and A20-OFP when injected subcutaneously (n=3, two-way ANOVA with Šídák’s multiple comparisons test). **b,** Kaplan-Meier survival curves of mice injected with 1.0 or 0.1 million A20 WT, or 1 million A20-OFP (each individual condition is analyzed against the 1 million A20 WT group using the Mantel-Cox test). **c,** Representative images of livers and spleens harvested from naïve mice or systemic A20-bearing mice harvested on day 25 or at the terminal stages. **d,** Percentage of A20 OFP^+^ tumor cells in livers on Day 22 after 1 million A20-OFP cells were injected retro-orbitally (n=2-3, unpaired t test). **e,** Kaplan-Meier survival analysis of mice bearing systemic A20 tumor cells treated with A20 neoantigen vaccination plus dual immune checkpoint therapy with or without depletion of CD8□ or CD4□ T cells (each individual condition is analyzed against the no depletion group using the Mantel-Cox test).

**Supplementary Figure 4:**
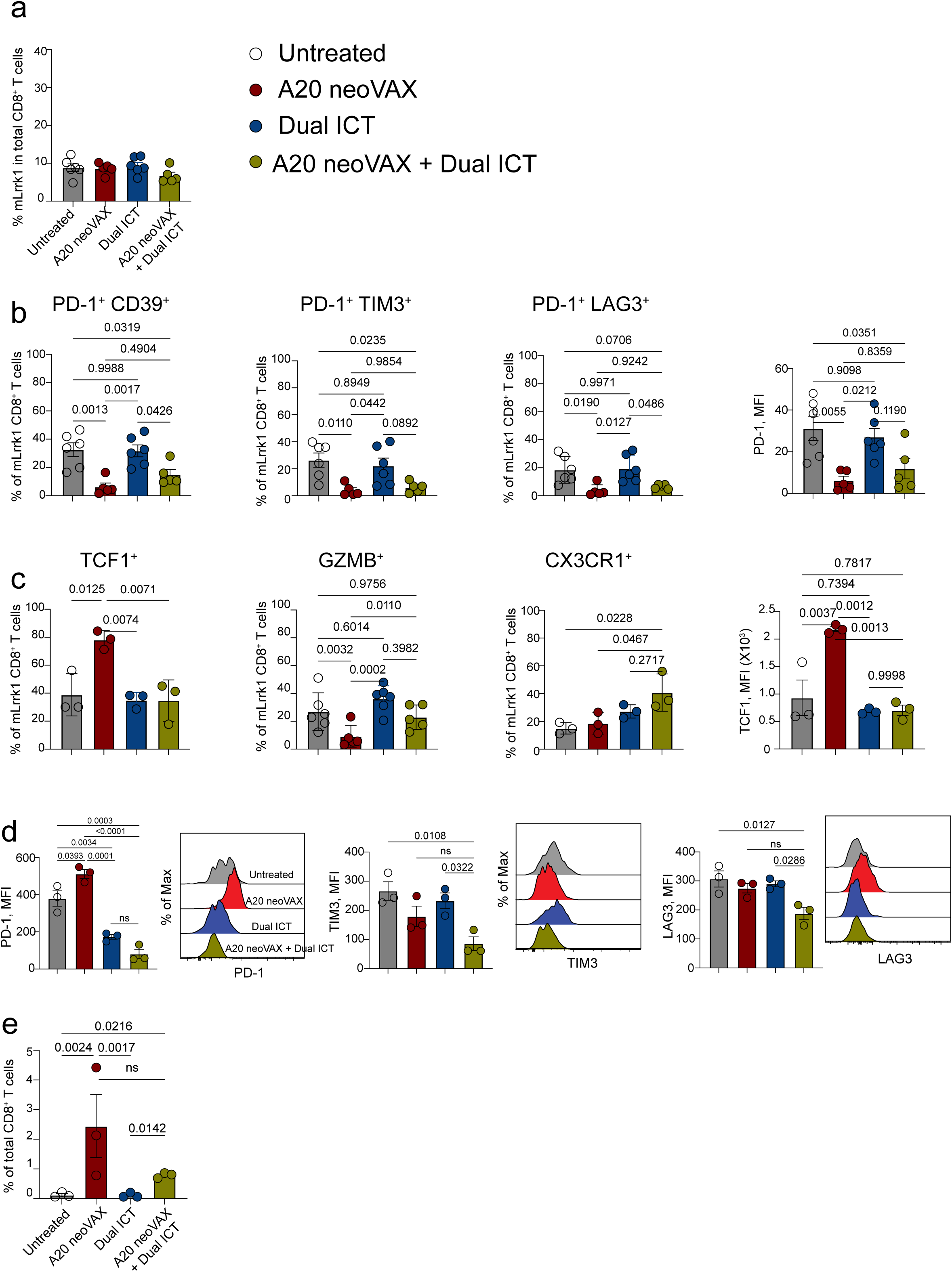
The addition of dual ICT in combinatorial therapy induced early skewing of vaccine-induced stem-like tumor-specific CD8^+^ T cells toward an effector-associated differentiation program. **a,** Frequencies of mLrrk1-specific CD8□ T cells among total CD8□ T cells in the liver of mice from the indicated treatment groups assessed 10 days after tumor inoculation. **b-c**, Livers were harvested from treated mice 10 days after systemic A20 tumor inoculation, followed by comprehensive flow cytometric analysis of mLrrk1-specific CD8□ T cells. **b,** Frequency of mLrrk1-specific CD8□ T cells coexpressing PD-1 and CD39, PD-1 and TIM-3, or PD-1 and LAG-3 or mean fluorescence intensity of PD-1 expression. **c,** Frequency of TCF1 (stem-like), GZMB (cytotoxic), or CX3CR1 (effector) in mLrrk1-specific CD8□ T cells. **d,** Mean fluorescence intensity for the expression of PD-1, TIM-3, and LAG-3 in mLrrk1-specific CD8□ T cells in the liver 15 days post tumor injection. **e,** Frequency of IFNγ and TNFα expressing CD8 TILs, isolated from the liver 15 days post tumor injection, following ex vivo stimulation with the indicated peptides for 4 h in the presence of Golgi Plug, followed by intracellular cytokine staining. Data are shown as mean□±□ s.e.m., with each symbol representing one mouse. Statistical analysis was performed using one-way ANOVA with Tukey’s multiple comparisons test.

**Supplementary Figure 5:**
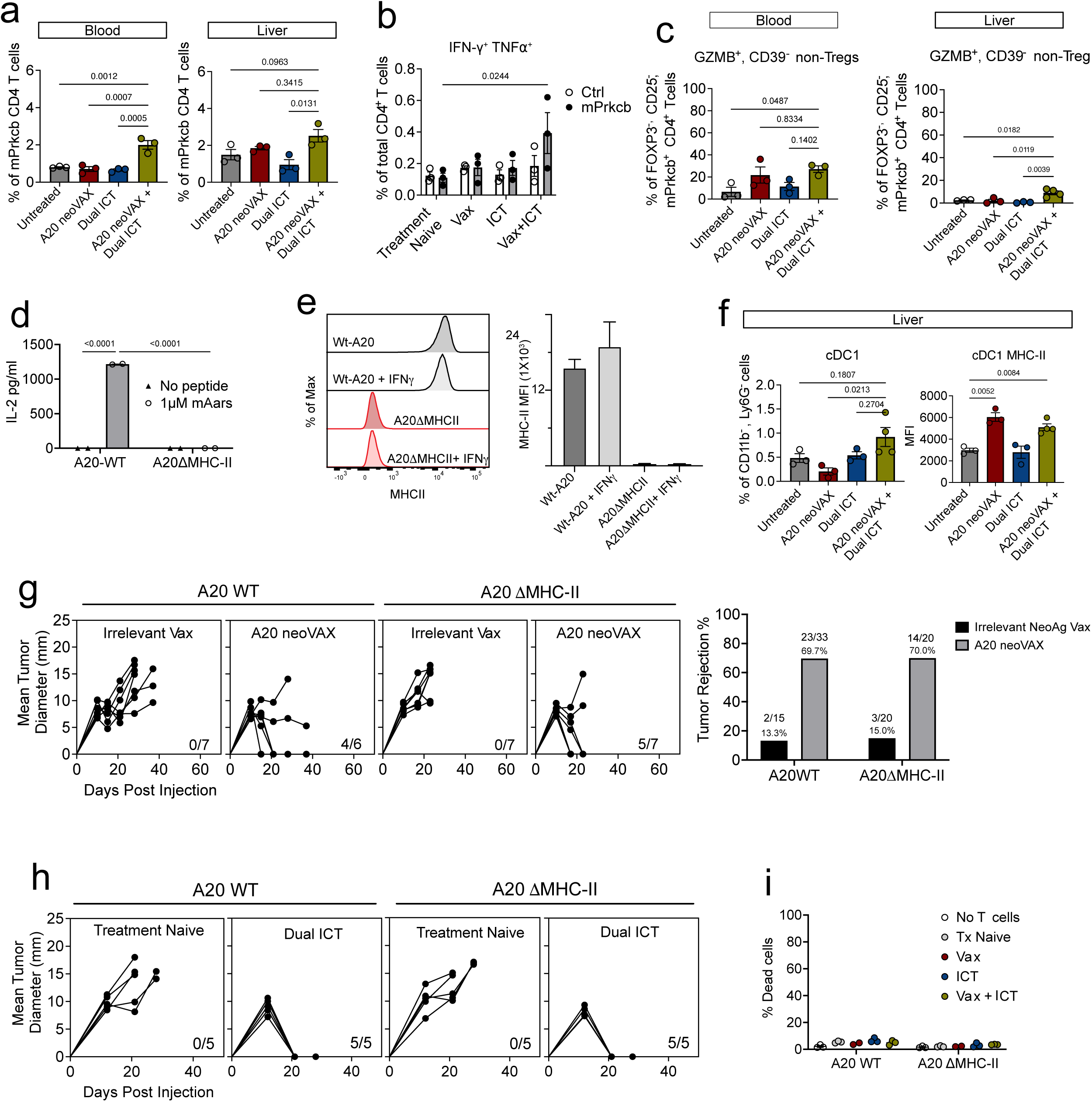
A20 neoVAX and dual ICT synergistically induced cytotoxic CD4^+^ TILs, but MHC-II expression on A20 cells was not required for efficient immunotherapies in both subcutaneous and systemic A20 models. **a,** Frequency of mPrkcb-specific CD4□ T cells among total CD4□ T cells in blood and liver assessed 10 days post tumor injection across different treatment groups. (**b, left)** Frequency of IFNγ and TNFα expressing CD4 TILs, isolated from the liver 15 days post tumor injection, following ex vivo stimulation with 1 γMof mPrkcb or control peptides for 4 h in the presence of Golgi Plug, followed by intracellular cytokine staining. **c,** Frequency of cytotoxic CD4^+^ T cells in blood and liver 15 days post tumor injection. **d,** IL-2 production from mAars T cell hybridoma using A20WT or A20ΔMHC-II as antigen-presenting cells (n=2 for each condition, two-way ANOVA with Tukey’s multiple comparisons test). **e,** Confirmation of MHC-II knockdown in A20 cells. **f,** Frequency of cDC1 and their MHC-II expression in the tumor and liver 15 days post tumor injection. For a–d, unless otherwise noted, data are shown as mean □±□ s.e.m., with each symbol representing one mouse. Statistical analysis was performed using one-way ANOVA with Tukey’s multiple comparisons test. **g,** Representative growth curves **(left)** of subcutaneous tumors established by A20WT or A20ΔMHC-II with treatment of irrelevant cancer vaccine or A20 neoantigen vaccine and summary **(right)** of repeated experiments. **h,** Representative growth curves of subcutaneous tumors established by A20WT or A20ΔMHC-II with or without dual ICT treatment. **(i)** Frequency of dead A20 tumor cells expressing NIR following co-culture with CD4□ TILs harvested from mice treated with multiple regimens as indicated, comparing wt-A20 and A20 ΔMHC-II targets (n=3 for each condition, two-way ANOVA with Tukey’s multiple comparisons test).

**Supplementary Table 1.**
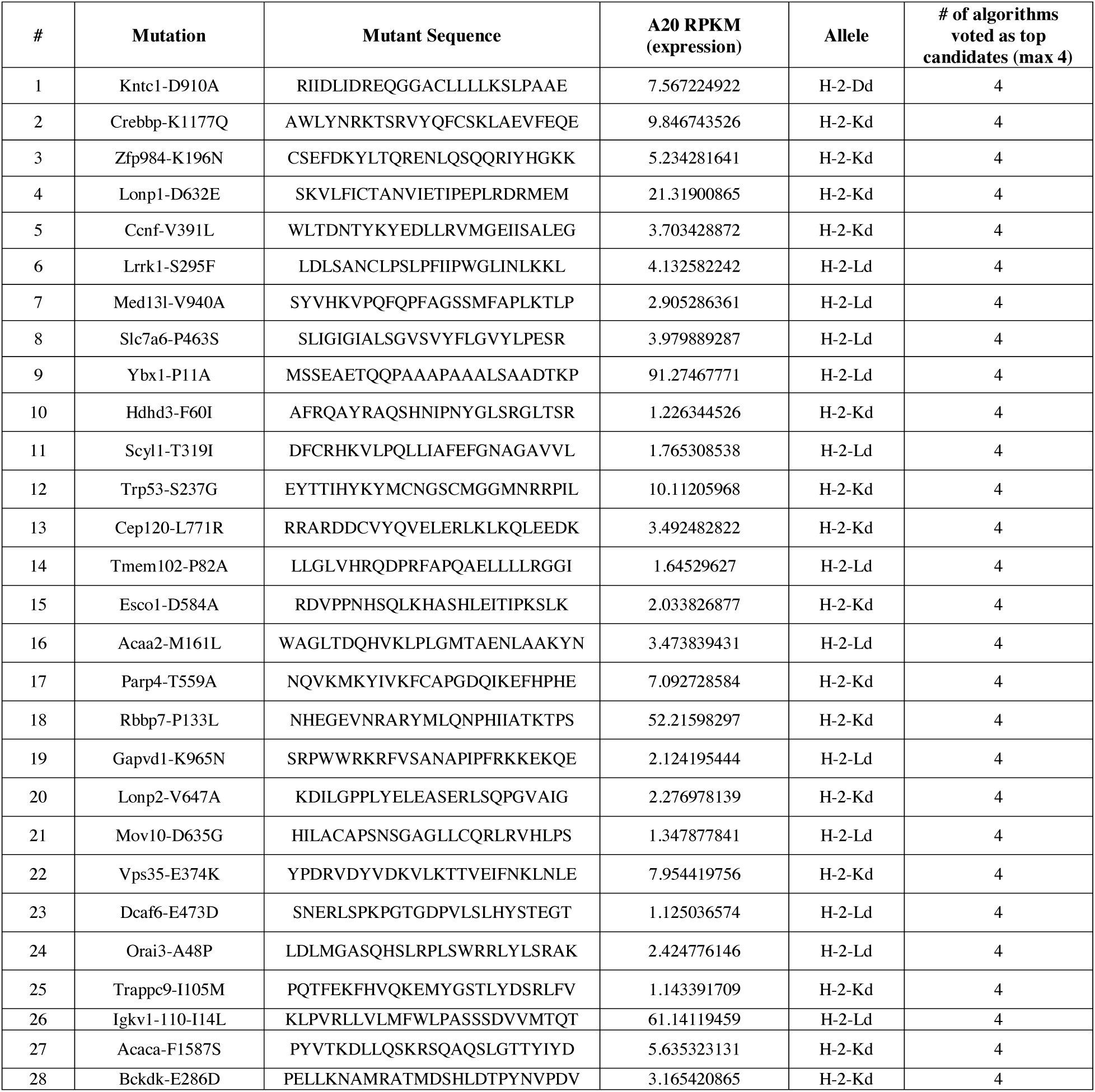

**Supplementary Table 2.**
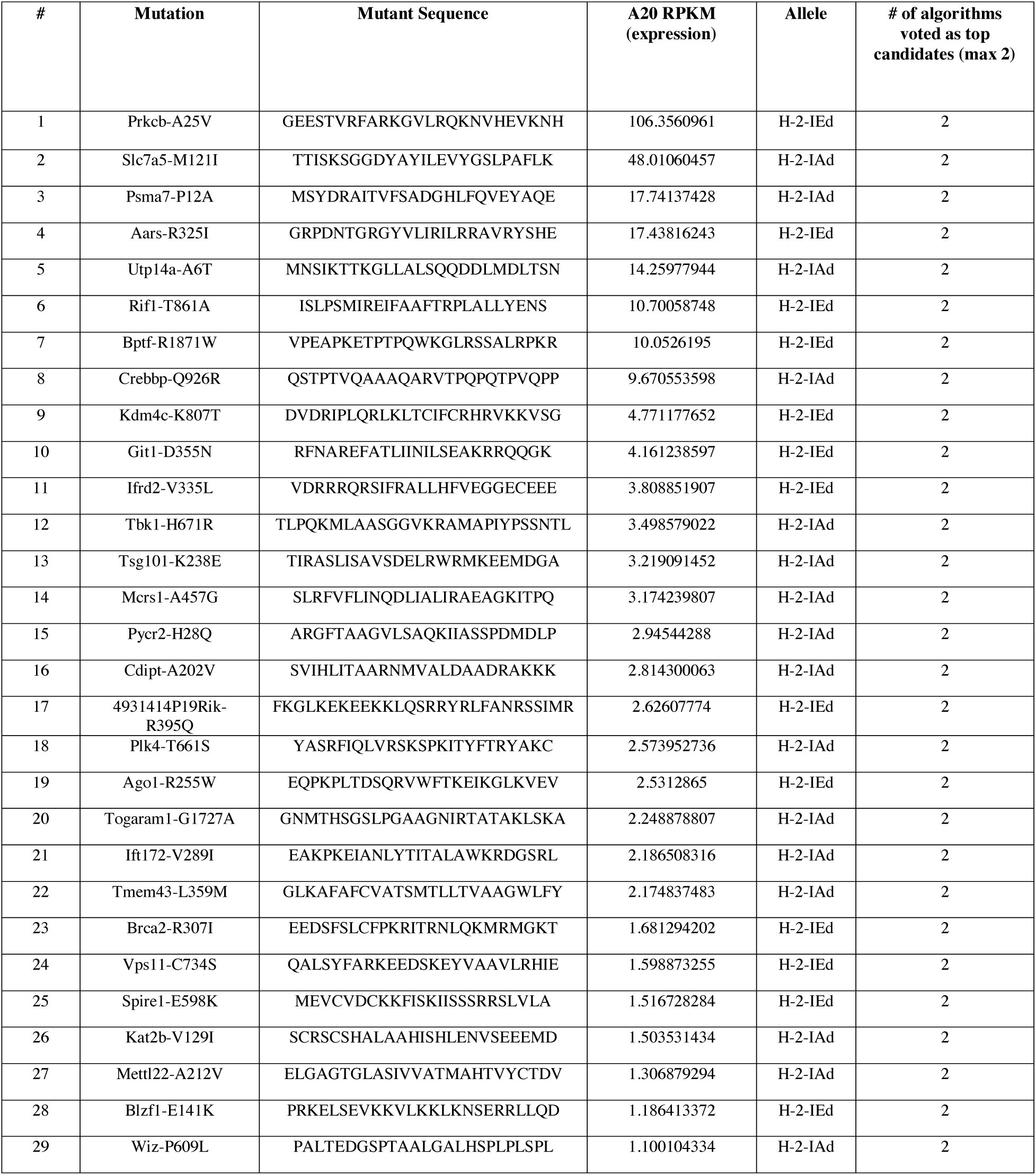

## Materials and Methods

### Mice

Female wild-type BALB/c mice (Charles River, strain code 028) were purchased from Charles River for experiments involving A20 tumor cells. BALB/c RAG2^−/−^ mice were bred and maintained in-house by our group. All in vivo experiments were performed in a specific-pathogen-free facility using mice between 8 and 12 weeks of age. All animal housing, feeding, and experiments were approved and performed in accordance with procedures approved by the AAALAC-accredited Animal Studies Committee of Washington University in St. Louis and followed all relevant ethical regulations.

### Culture and transplantation of tumor cell lines

A20 cells were obtained from Dr. Ronald Levy’s lab (Stanford University). Tumor cells were cultured in Roswell Park Memorial Institute 1640 medium (RPMI; Cytiva, Cat# SH30096.02) supplemented with 10% fetal calf serum (FCS; Cytiva, Cat# SH30071.03). All cell lines used in this study were passaged 3–4 times before use and were confirmed to be free of mycoplasma contamination. Briefly, 1 mL of antibiotic-free cell culture supernatant was submitted to IDEXX Laboratories (Columbia, MO, USA) for mycoplasma testing, and all results were negative. For injection, A20 cells were washed 3 times with PBS (Cytiva, Cat# SH30028.03) and resuspended at a density of 6.67×10^6^ cells per mL in PBS.

#### Subcutaneous tumor transplantation

For the subcutaneous model, a total of 1×10^6^ cells in 150 γL were injected subcutaneously into the right rear flanks of syngeneic recipient BALB/c mice. Following tumor transplantation, animals were randomly assigned to treatment groups. Tumor growth was measured by caliper measurement, and individual growth curves are represented as the average of two perpendicular diameters. Tumor-bearing animals were euthanized when the maximal tumor diameter reached 20 mm in one direction.

#### Systemic tumor transplantation

A20-OFP and A20 cells were cultured with the same protocol as described above. For injection, tumor cells were washed 3 times with PBS (Cytiva, Cat# SH30028.03) and resuspended at a density of 6.67×10^6^ cells per mL in PBS. A total of 1×10^6^ cells in 150 γL was injected intravenously via the tail vein. Tumor-bearing animals were euthanized when the mice developed hepatomegaly (mice’s weight increased by over 20%, caused by tumor burden) or other symptoms (hunched, moribund, paralyzed, etc.).

### RNA sequencing and whole-exome sequencing

DNA and RNA were isolated from in vitro cultured A20 cells using QIAamp DNA Kits (Qiagen, Cat# 56304) and RNeasy Kits (Qiagen, Cat# 74104) according to the manufacturer’s instructions. DNA was isolated from the tails of female BALB/c mice to serve as the control. DNA and RNA (N = 1 per condition) were sent to the Genomic Services Laboratory at the Steve and Cindy Rasmussen Institute for Genomic Medicine at Nationwide Children’s Hospital in Columbus, Ohio, for whole exome and RNA sequencing (RNA-seq). Whole exome and bulk RNA-seq were performed as previously described^21,44,77^.1–3 Briefly, exome-sequencing libraries were generated using the NEBNext Ultra II FS Kit (NEB, Cat# E7805), and paired-end 151 base pair reads were sequenced on the NovaSeq6000 to aim for 100x coverage in normal and 250x in tumor samples. Tumor RNA was subjected to DNase treatment and ribodepletion before library construction using the NEBNext Ultra II Directional RNA library prep kit for Illumina (NEB, Cat# E7760). Paired-end 151 base pair reads were generated on the NovaSeq6000 to generate a minimum of 80 million reads per sample. Raw sequencing reads were processed using the publicly available Common Workflow Language (CWL) pipelines from the McDonnell Genome Institute/Griffith Lab analysis-workflows repository (https://github.com/genome/analysis-workflows), executed via Cromwel. Tumor and normal exome reads were processed with the ’somatic_exome_nonhuman.cwl’ pipeline: reads were aligned to the mouse reference genome (GRCm38) using BWA-MEM, and somatic variants were called with a multiple variant callers, merged into a combined call set, annotated with the Ensembl Variant Effect Predictor (VEP), and filtered to remove likely germline and artifactual calls. Tumor RNA-seq reads were processed with the ’rnaseq.cwl’ pipeline: reads were adapter-trimmed and aligned to GRCm38 using the splice-aware aligner HISAT2, with gene- and transcript-level expression quantified using StringTie and Kallisto, respectively.

### MHC-I and MHC-II Neoantigen prediction of A20

Neoantigen predictions were performed using mouse A20 exome data, as previously described^21,44,77^. The VCF file of somatic variants from above was processed with the ’pvacseq.cwl’ subworkflow, which annotates variants with RNA-seq expression and read-count/coverage support and runs pVACseq for neoantigen prediction and ranking. Peptide sequences arising from each somatic mutation were evaluated for lengths 8–11 using eight Class I prediction algorithms in pVACseq and four class I prediction algorithms by hmMHC. Class II prediction was performed using four algorithms supported by pVACseq and two algorithms by hmMHC that were informed by Mass Spec data of many MHC-II binding self-peptides and neoantigens that have been compiled by the Washington University community and others. Potential neoantigens were filtered and prioritized according to the following criteria: the number of algorithms with good binding predictions for the mutant epitopes, binding predictions for the wildtype epitopes, variant DNA/RNA VAF, variant DNA/RNA coverage, and variant transcript expression.

### Peptides and neoantigens vaccination protocol

For in vivo vaccination, [30 μg of mLrrk SLP (Seq; LDLSANCLPSLPFIIPWGLINLKKL (MHC-I A20 neoVAX), 1.5 ng of mPrkcb (Seq; GEESTVRFARKGVLRQKNVHEVKNH) and/or mAars (Seq; GRPDNTGRGYVLIRILRRAVRYSHE) (MHC-II A20 neoVAX) or 30 μg of mLrrk SLP plus 1.5 ng of mPrkcb and/or mAars (A20 neoVAX)] were mixed with 50 μg of polyinosinic–polycytidylic acid complexed with poly-L-Lysine and carboxymethylcellulose (poly-ICLC, Oncovir, Inc) diluted in PBS. For therapeutic vaccination, tumor-bearing mice were injected subcutaneously with the vaccines (150 μl) on days 6 and 17 post-tumor transplant. In experiments using the combination of vaccination and ICT, vaccines were administered subcutaneously as indicated on the figures. The mTnpo3 SLP (Sequence; YGMEEGCRQGLSYMLQALCIPTFQL) was used as an irrelevant antigen^78^. All 25-mer peptides used for neoantigen screening were purchased from Genscript with >90% purity (Supplementary Tables 1 and 2). All 25-mer SLP neoantigen peptides used for vaccination and in vivo experiments were purchased from Genscript or Peptide 2.0 and purified by high-performance liquid chromatography (HPLC) to >95% purity. For all the mutant peptides, the mutant amino acid was placed in the center of the peptide with wild-type 12-aa flanks on both sides. The identity of the peptide was also validated by mass spectrometry.

### In vivo antibody treatment

For immune checkpoint therapy, rat IgG2a anti-PD-1 (RMP1-14, Leinco Technologies, Product No.: P372), mouse IgG2b anti-CTLA4 (9D9, Leinco Technologies, Product No.: C2856), and anti-PD-L1 (10F.9G2, Leinco Technologies, Product No.: P371) antibodies were used. Rat IgG2a isotype control antibody (1-1, Leinco Technologies, Product No.: R1367) was also used in some experiments as a control treatment. Mice were injected intraperitoneally with 200 γg of each antibody on days 6, 9, and 12 after tumor transplantation for subcutaneous A20 tumors or on days 3, 6, 9, and 12 for systemic A20 tumors.

Anti-CD8 (53-6.7, Leinco Technologies, Product No.: C2848) and anti-CD4 (GK1.5, Leinco Technologies, Product No.: C2838) mAbs were used for CD8 and CD4 T cell depletion, respectively. Mice were injected intraperitoneally with 150 γg of each antibody every 3 days, starting 1 day before tumor inoculation.

### CD8-IL2 and CD8-IL21 administration

Mouse CD8-IL2 and mouse CD8-IL21 were generated and kindly provided by Asher Biotherapeutics and previously described^55,61^. For the current study, CD8-IL2 and CD8-IL21 were administered one time at a dose of 1 mg/kg diluted in PBS and injected intraperitoneally on day 12 post-tumor inoculation.

### Peptide-MHC Tetramer Staining and Flow Cytometry

Tetramer staining for mLrrk1-specific CD8^+^ T cells and mPrkcb-specific CD4^+^ T cells was performed as previously described. Biotinylated peptide-MHC monomers were obtained from the Bursky Center Immunomonitoring Laboratory at Washington University School of Medicine in St. Louis and conjugated with fluorophore-labeled streptavidin. Samples were incubated with tetramers at 37□ °C for 30□ min before surface molecule staining.

For multi-colour flow cytometry, the following antibodies were used: [BioLegend: CD45 (30-F11;1:500; cat# 103138); BD Bioscience: THY1.2 (53-2.1; 1:500; cat# 565257), CD4 (RM4-5: 1:500; cat# 612843) and CD8 (53.6.7; 1:200; cat# 564920)]. Zombie NIR (BioLegend) was used to stain for cellular viability. The BD Cytofix/Cytoperm Plus kit (BD Biosciences) was used according to the manufacturer’s protocol for intracellular staining of cytokines and other intracellular proteins.

### ELISPOT

For neoantigen candidate screening, cells from tumors were enriched for CD8^+^ or CD4^+^ T cells using the Miltenyi mouse CD8^+^ or CD4^+^ enrichment kits. 500,000 splenocytes from RAG2^-/-^mice were pulsed with 1 or 10 γM 25-mer peptides and were used to stimulate enriched T cells overnight at 37□ in anti-mouse IFNγ-coated ELISPOT plates (Immunospot). 25 µg/ml phytohemagglutinin (PHA) was used as a positive control. For SLP neoantigen immunogenicity analysis, splenocytes from vaccinated mice were harvested and pulsed with 10 γM 25-mer peptides. Plates were developed according to the manufacturer’s protocol. The results were quantified by a CTL ImmunoSpot S6 Universal machine and Professional 6.0.0 software.

### In vivo cytotoxicity assay

Splenocytes were collected from naïve BALB/c mice and used as target cells. Target cells were pulsed with 1γM mLrrk1 or control peptide overnight at 37□. After incubation, peptide-pulsed cells were stained with 5γM or 0.5γM CellTrace™ CFSE (Thermo Fisher Scientific) as CFSE^high^ (mLrrk1 peptide pulsed) or CFSE^low^ (control peptide pulsed), respectively. Stained cells were washed three times and combined at a 1:1 ratio in PBS. 20×10^6^ cells were injected retro-orbitally into tumor-bearing mice 13 days post-tumor injection, with or without dual ICT treatment. Non-tumor-bearing mice were used as controls. After 24 hours of target cell transfer, splenocytes from the transferred mice were harvested and stained with Zombie NIR viability dye (BioLegend).

Live CFSE-positive cells were quantified to calculate the relative lysis of mLrrk1 CFSE^high^ targets compared to control CTV^low^ targets in tumor-naïve, treatment-naïve tumor-bearing mice and ICT-treated tumor-bearing mice. mLrrk1-specific killing was represented as [1 − (tumor-naive control CFSE low vs high ratio/tumor-bearing experimental CFSE low vs high ratio)] × 100%.

### In vitro cytotoxicity assay

CD8□ T cells were isolated from tumors using the Miltenyi mouse CD8□ T cell enrichment kit (Miltenyi Biotec Cat# 130-104-075) according to the manufacturer’s protocol. Enriched CD8□ TILs were stained with PE-conjugated tetramer, prepared as previously described, and detected using PE-conjugated streptavidin (BioLegend Cat# 405204). Tetramer-specific T cells were enriched using anti-PE microbeads (Miltenyi Biotec Cat# 130-048-801) according to the manufacturer’s protocol. Enriched tetramer–specific CD8□ T cells were co-cultured with WT A20 or A20 ΔMHC-I target cells at 1:1 effector-to-target ratios. After 24 hours, cells were washed twice with PBS and stained with [BioLegend: CD45 (30-F11;1:500; cat# 103138) and CD19 (ID3; 1:500; cat #152410)]. Zombie NIR fixable viability dye (BioLegend Cat# 423105) was used to determine the frequency of dead Zombie NIR□ CD45^+^CD19□ tumor cells.

### Hybridoma generations

Bulk CD4□ T cells were isolated from the spleens of SLP-vaccinated mice after two doses of vaccine treatment and stimulated in vitro with peptide-pulsed (1µM) naive splenocytes. Activated CD4□ T cells were fused with BW5147 thymoma cells to generate hybridomas, which were cloned by limiting dilution. To determine antigen specificity, splenocytes from naïve mice were pulsed with peptide (0-10 µM) and co-cultured overnight with hybridoma cells (5×10^4^ hybridomas with 5×10^4^ peptide-pulsed splenocytes). Supernatants were collected, and IL-2 production was quantified by ELISA.

### CODEX Imaging

BALB/c mice bearing A20 tumors were treated with either Control Ab or α-CTLA4 + α-PD-1, and tumors were harvested 13 days later, fresh frozen, cut into 8 γm-thick tissue slices, and stained with the CODEX antibody panel as described^79^. CD19 is used as a marker for A20 lymphoma cells.

### NeoAg vaccinations

For therapeutic NeoAg vaccinations, the indicated peptides (MHC-I SLPs: 30 µg; MHC-II SLPs: 15 µg unless otherwise annotated) were mixed with 25 γg of polyinosinic–polycytidylic acid complexed with poly-l-lysine and carboxymethylcellulose (poly-ICLC; Oncovir) diluted in PBS^44^ Tumor-bearing mice were injected intravenously or subcutaneously with the vaccines in 150 γl PBS on days 3, 10, and 17 post-tumor inoculation.

### Adoptive transfer experiments

Wild-type tumor-bearing mice were vaccinated with the A20 NeoAg vaccine on days 3 and 10 post-tumor transplantation. Splenocytes from vaccinated mice were harvested 15 days after tumor injections and enriched for total T cells (THY1.2 beads), CD8^+^ T cells, or CD4^+^ T cells using the Miltenyi cell enrichment kits. Total T cells (5-10×10^6^ ) and either CD8 or CD4 T cells were transferred into Rag2^−/−^ mice that were injected with 2.5□ ×□ 10^5^ tumor cells 3 days prior to T cell transfer.

### Generation of A20-OFP cell line

293T cells were used to produce OFP-encoding lentivirus according to the standard protocol (TransIT® Lentivirus System). 1-2×10^6^ A20 cells were seeded in 500 µl virus supernatant in a 12-well plate. Polybrene was added to the supernatant at a final concentration of 8 µg/ml. The cells were spinoculated in a 12-well plate at 700g, 20-25□ for 1 hour. Cells were cultured in virus supernatant overnight. Cells were cultured in R10 media and sorted for OFP^+^ cells.

### Ex vivo fluorescence imaging

Ex vivo fluorescence imaging of tumor-infiltrated organs was performed using the IVIS SpectrumCT imaging system (PerkinElmer) at the Washington University Molecular Imaging Center. Following tissue collection, organs were imaged under identical acquisition settings for all experimental groups. Fluorescence intensity was quantified using Living Image software (version 4.8; PerkinElmer) by defining regions of interest (ROIs) encompassing each organ. Total fluorescence was measured and used to quantitatively compare experimental groups.

### Flow cytometry

Flow antibodies were purchased from: [BioLegend: CD45 (30-F11;1:500; cat# 103138), IFN-γ (XMG1.2; 1:100; cat# 505808), TNF-α (XMG1.2; 1:100; cat# 505826), PD-1 (29F.1A12; 1:200; cat# 135228), GZMB (QA16A02; 1:50; cat# 372216), TIM-3 (RMT3-23; 1:200; cat# 119723), CD25 (PC61; 1:100; cat# 102036), CD154 (MR1;1:100; cat# 106506), CD152 (UC10-4B9; 1:100; cat# 106318), IL-2 (Jes6-5H4: 1:50; cat# 503826), LILRB4 (H1.1: 1:200; cat# 144904), CD11b (M1/70: 1:800; cat# 101226), XCR1 (ZET: 1:100; cat# 148206), MHC-II (M5/114.15.2: 1:1000; cat# 107641), CD11c (N418: 1:200; cat# 117336), CD172a (P84: 1:500; cat# 144008), (Zombie NIR fixable viability dye; 1:500; cat# 423106), CD86 (GL-1; 1:200; cat# 105042), CD80 (16-10A1; 1:200; cat# 104712), CD40 (FGK45; 1:100; cat# 157506), CD64 (X54-5/7.1; 1:100; cat# 139304), T-BET (4B10; 1:100; cat# 644816), KLRG1 (2F1; 1:100; cat# 138421), B220 (RA3-6B2; 1:100; cat #103206), CD19 (ID3; 1:500; cat #152410), CX3CR1 (SA011F11; 1:100; cat# 149031); BD Bioscience: CD39 (Y23-1185; 1:400; cat# 567105), THY1.2 (53-2.1; 1:500; cat# 565257), LY6G (1A8; 1:500; cat# 563979), CD278 (C398.4A, 1:100; cat# 568249), F4/80 (T45-2342; 1:100; 565614), CD223 (T47-530; 1:100; cat# 744727), Ki-67 (B56; 1:100; cat# 561284), CD4 (RM4-5: 1:500; cat# 612843) and CD8 (53.6.7; 1:200; cat# 564920); or eBioSciences: FOXP3 (Fjk-16a; 1:50; cat# 11-5773-82) and iNOS (CXNFT; 1:100; cat# 12-5920-82)]. Foxp3/Transcription factor Staining kit (ThermoFisher; cat# 00-5523-00) was used to stain Foxp3 and other intracellular proteins according to the manufacturer’s protocol. BD FACSDIVA software V9.1 on Symphony 3, LSRFortessaX20, or ARIA-II was used for collecting the flow cytometry data and for sorting, respectively. Data were analyzed using FlowJo software version 10.10.

Zombie NIR (BioLegend) was used to stain for cellular viability. The BD Cytofix/Cytoperm Plus kit (BD Biosciences) was used according to the manufacturer’s protocol for intracellular staining of cytokines and other intracellular proteins. Foxp3/Transcription factor Staining kit (00-5523-00, Thermo Fisher) was used to stain FOXP3 and other intracellular proteins according to the manufacturer’s protocol.

Tetramer staining for mLrrk1-specific CD8^+^ T cells and mPrkcb-specific CD4^+^ T cells was performed as previously described^21,77^. Biotinylated peptide-MHC monomers were obtained from the Bursky Center Immunomonitoring Laboratory at Washington University School of Medicine in St. Louis and conjugated with fluorophore-labeled streptavidin. Samples were incubated with tetramers at 37□ °C for 30□ min before surface molecule staining.

### Multiplex cytokine assay

Cells from livers were sorted for mLrrk1-specific CD8^+^ T cells and mPrkcb CD4^+^ T cells from tumor-bearing mice receiving different treatments. Sorted cells (20,000 cells per sample) were stimulated in a serum-free medium with 10^6^ irradiated splenocytes (isolated from naive mice) pulsed with 1-3µM mLrrk1 or mPrkcb SLP. Following a 72-h incubation, secretion of multiple cytokines was measured using a flow-based customized ProcartaPlex cytokine panel (Luminex Technologies) following the manufacturer’s protocol.

### Intracellular cytokine assay

Splenocytes (10^5^) harvested from naïve mice were irradiated (30 Gy) and pulsed with 1 γg/ml of the relevant peptide, and TILs (1-2 × 10^6^) were subsequently added, and the cell suspension was incubated at 37 °C. GolgiPlug (BD Bioscience) was added 1 hr later and incubated for another 4 hr. Cells were stained for different surface markers, including the live/dead marker (NIR), then permeabilized using the intracellular permeabilization kit (BD), followed by staining for [BioLegend: IFN-γ (XMG1.2; 1:100; cat# 505808), TNF-α (XMG1.2; 1:100; cat# 505826), IL-2 (Jes6-5H4: 1:50; cat# 503826)].

### Generation of A20ΔMHC-I and A20ΔMHC-II cell lines

A20ΔMHC-II cell lines were generated by CRISPR-Cas9 using a previously reported protocol^80^. The approach used the PX330-based plasmid (obtained from Addgene) encoding an sgRNA scaffold, SpCas9, and mCherry. Guide sequences (GGTGACATTGACGACTGGAG and GGAGAGTACAGTCACCTCTG for H2-Ea; CAGATACATCTACAACCGGG and GTGCGCTACGACAGCGACGT for H2-Ab1) were designed to target the H2-Ea and H2-Ab genes, ordered from IDT, and incorporated into separate plasmids. A20ΔMHC-I cell lines were generated using a similar approach, with guide sequences (GTATTACAGGGCCTACCTAG and TGACATCAACTTGAGATCTG For H2-Kd; TACTGGCAGTTCGCCTACGA and CTGGTTGTAGTAGCGCAGCG for H2-Dd; GGTGACTTCACCTTTAGATC and TGTCGGCTATGTGGACAACA for H2-Ld). Engineered plasmids were electroporated into A20 cells using Lonza SF Cell Line 4D-Nucleofector™ X Kit L V4XC-2024. Cells were first sorted to enrich for mCherry+ cells. After expansion, cells were sorted twice more to enrich for mCherry^-^ MHC-II^-^ cells. A control plasmid with scrambled sgRNA sequences (GCTTAGTTACGCGTGGACGA) was used to generate control A20 cells (A20-SCR), which served as A20WT in some experiments.

IL-2 ELISA was used to analyze the supernatant of mAars-specific hybridoma cells cocultured with A20WT cells or A20ΔMHC-II cells. A20WT cells or A20ΔMHC-II cells were pulsed with 1 M mAars SLPs for 1 hour at 37□ and washed with PBS. Tumor cells were then cocultured with mAars-specific hybridoma at a 1:1 ratio for 12 hours, and the supernatant was analyzed by IL-2 ELISA following the manufacturer’s protocol (BioLegend).

### Statistics

Statistical analyses were performed using GraphPad Prism v10.4.1 (GraphPad Software). Statistical comparisons were conducted using unpaired, two-tailed Student’s t tests, one-way ANOVA, or two-way ANOVA, as appropriate. Detailed information regarding the statistical test used for each experiment is provided in the corresponding figure legends. We selected group sizes based on prior experience with these tumor models and published studies, and we did not perform a power calculation to determine sample size.

## Acknowledgements

We thank all members of the Schreiber laboratory for discussions and technical support; K. Link, L. Yang, and D. Bender of the Immunomonitoring Laboratory (IML), who provided tetramers for MHC-I and MHC-II neoantigens and the cytokine ELISPOT assessments; and J. Prior of the WashU Molecular Imaging Center (MIC), who provided technical support in ex vivo fluorescent imaging.

## Funding Information

This work was supported by grants to RDS from the National Cancer Institute of the US National Institutes of Health (1R21CA313364-01), and the Parker Institute for Cancer Immunotherapy. HS was supported by the National Cancer Institute of the US National Institutes of Health (1R21CA313364-01) and a Young Investigator Award from the Cancer Research Foundation. TAF is supported by Blood Cancer United (Specialized Center of Research), the Paula and Rodger Riney Foundation, and the Steinberg Family Cancer Research Fund. RDS, TAF, MG and OG are supported by a Blood Cancer United SCOR grant. The IML is supported by the Andrew M. and Jane N. Bursky Center for Human Immunology and Immunotherapy Programs and the Alvin J. Siteman Comprehensive Cancer Center. The latter is supported by a National Cancer Institute of the US National Institutes of Health Cancer Center support grant (P30CA91842).

## Conflicts of Interest

R.D.S. is a cofounder, scientific advisory board member, stockholder, and royalty recipient of AsherBio and is a scientific advisory board member for Revity Inc. T.A.F. holds equity in Orca Bio, Indapta Therapeutics, and Wugen, consults for Wugen, and received research funding from Miltenyi and AI Proteins. All other authors declare no conflicts of interest.

## Notes

### Competing Interest Statement

The authors have declared no competing interest.

